# Quasiperiodic rhythms of the inferior olive

**DOI:** 10.1101/408112

**Authors:** Mario Negrello, Pascal Warnaar, Vincenzo Romano, Cullen B. Owens, Sander Lindeman, Elisabetta Iavarone, Jochen K. Spanke, Laurens W.J. Bosman, Chris I. De Zeeuw

## Abstract

Inferior olivary activity causes both short-term and long-term changes in cerebellar output underlying motor performance and motor learning. Many of its neurons engage in coherent subthreshold oscillations and are extensively coupled via gap junctions. Studies in reduced preparations suggest that these properties promote rhythmic, synchronized output. However, how these properties interact with synaptic inputs controlling inferior olivary output in intact, awake behaving animals is poorly understood. Here we combine electrophysiological recordings in awake mice with a novel and realistic tissue-scale computational model of the inferior olive to study the relative impact of intrinsic and extrinsic mechanisms governing its activity. Our data and model suggest that if subthreshold oscillations are present in the awake state, the period of these oscillations will be transient and variable. Accordingly, by using different temporal patterns of sensory stimulation, we found that complex spike rhythmicity was readily evoked but limited to short intervals of no more than a few hundred milliseconds and that the periodicity of this rhythmic activity was not fixed but dynamically related to the synaptic input to the inferior olive as well as to motor output. In contrast, in the long-term the average olivary spiking activity was not affected by the strength and duration of the sensory stimulation, while the level of gap junctional coupling determined the stiffness of the rhythmic activity in the olivary network during its dynamic response to sensory modulation. Thus, interactions between intrinsic properties and extrinsic inputs can explain the variations of spiking activity of olivary neurons, providing a conceptual framework for the creation of both the short-term and long-term changes in cerebellar output.

**AUTHOR SUMMARY:** Activity of the inferior olive, transmitted via climbing fibers to the cerebellum, regulates initiation and amplitude of movements, signals unexpected sensory feedback, and directs cerebellar learning. It is characterized by widespread subthreshold oscillations and synchronization promoted by strong electrotonic coupling. In brain slices, subthreshold oscillations gate which inputs can be transmitted by inferior olivary neurons and which will not - dependent on the phase of the oscillation. In our study, we tested whether the subthreshold oscillations had a measurable impact on temporal patterning of climbing fiber activity in intact, awake mice. We did so by recording neural activity of the postsynaptic Purkinje cells, in which complex spike firing faithfully represents climbing fiber activity. For short intervals (<300 ms), we found that many Purkinje cells indeed showed spontaneously rhythmic complex spike activity. However, our experiments designed to evoke resonant responses indicated that complex spikes are not predominantly predicated on stimulus history. Our realistic network model of the inferior olive explains the experimental observations via continuous phase modulations of the subthreshold oscillations under the influence of synaptic fluctuations. We conclude that complex spike activity emerges from a quasiperiodic rhythm that is stabilized by electrotonic coupling between its dendrites, yet dynamically influenced by the status of their synaptic inputs.

## INTRODUCTION

A multitude of behavioral studies leave little doubt that the olivo-cerebellar system organizes appropriate timing in motor behavior (1–3), perceptual function (4–6) and motor learning (7–10). Furthermore, the role of the inferior olive in motor function is evinced in (permanent and transient) clinical manifestations, such as tremors, resulting from olivary lesions and deficits (11–16). Although the consequences of olivary dysfunctions are rather clear, the network dynamics producing functional behavior are controversial. At the core of the controversy is the question whether inferior olive cells are oscillating during the awake state and whether these oscillations affect the timing of the inferior olivary output (17–19). The inferior olive is the sole source of the climbing fibers, the activity of which dictates complex spike firing by cerebellar Purkinje cells (for review, see (20)). Climbing fiber activity is essential for motor coordination, as it contributes to both initiation and learning of movements (8, 10, 21–26), and it may also be involved in sensory processing and regulating more cognitive tasks (27–30). Understanding the systemic consequences of inferior olivary spiking is therefore of great importance.

The dendritic spines of inferior olivary neurons are grouped in glomeruli, in which they are coupled by numerous gap junctions (10, 31–33), which broadcast the activity state of olivary neurons. Due to their specific set of conductances (34–40), the neurons of the inferior olive can produce subthreshold oscillations (STOs) (41–43). The occurrence of STOs does not require gap junctions per se (44), but the gap junctions appear to affect the amplitude of STOs and engage larger networks in synchronous oscillation (10,16,42). Both experimental and theoretical studies have demonstrated that STOs may mediate phase-dependent gating where the phase of the STO helps to determine whether excitatory input can or cannot evoke a spike (45, 46). Indeed, whole cell recordings of olivary neurons in the anesthetized preparation indicate that their STOs can contribute to the firing rhythm (42, 43) and extracellular recordings of Purkinje cells in the cerebellar cortex under anesthesia often show periods of complex spike firing around the typical olivary rhythm of 10 Hz (17, 47–49). However, several attempts to capture clues to these putative oscillations in the absence of anesthesia have, so far, returned empty handed (19, 50).

It has been shown that in the anesthetized state both the amplitude and phase of the STOs can be altered by synaptic inputs (10). Inhibitory inputs to the inferior olive originate in the cerebellar nuclei and have broadly distributed terminals onto compact sets of olivary cells (51–53). Excitatory terminals predominantly originate in the spinal cord and lower brainstem, mainly carrying sensory information, and in the nuclei of the meso-diencephalic junction in the higher brainstem, carrying higher-order input from the cerebral cortex (Fig. 1A) (15, 54, 55). In addition, the inferior olive receives modulating, depolarizing, level-setting inputs from areas like the raphe nuclei (55). Unlike most other brain regions, the inferior olive is virtually devoid of interneurons (56, 57). Thus, the long-range projections to the inferior olive in conjunction with presumed STOs and gap junctions jointly determine the activity pattern of the complex spikes in Purkinje cells. How these factors contribute to functional dynamics of the olive in awake mammals remains to be elucidated.

**Fig. 1.**
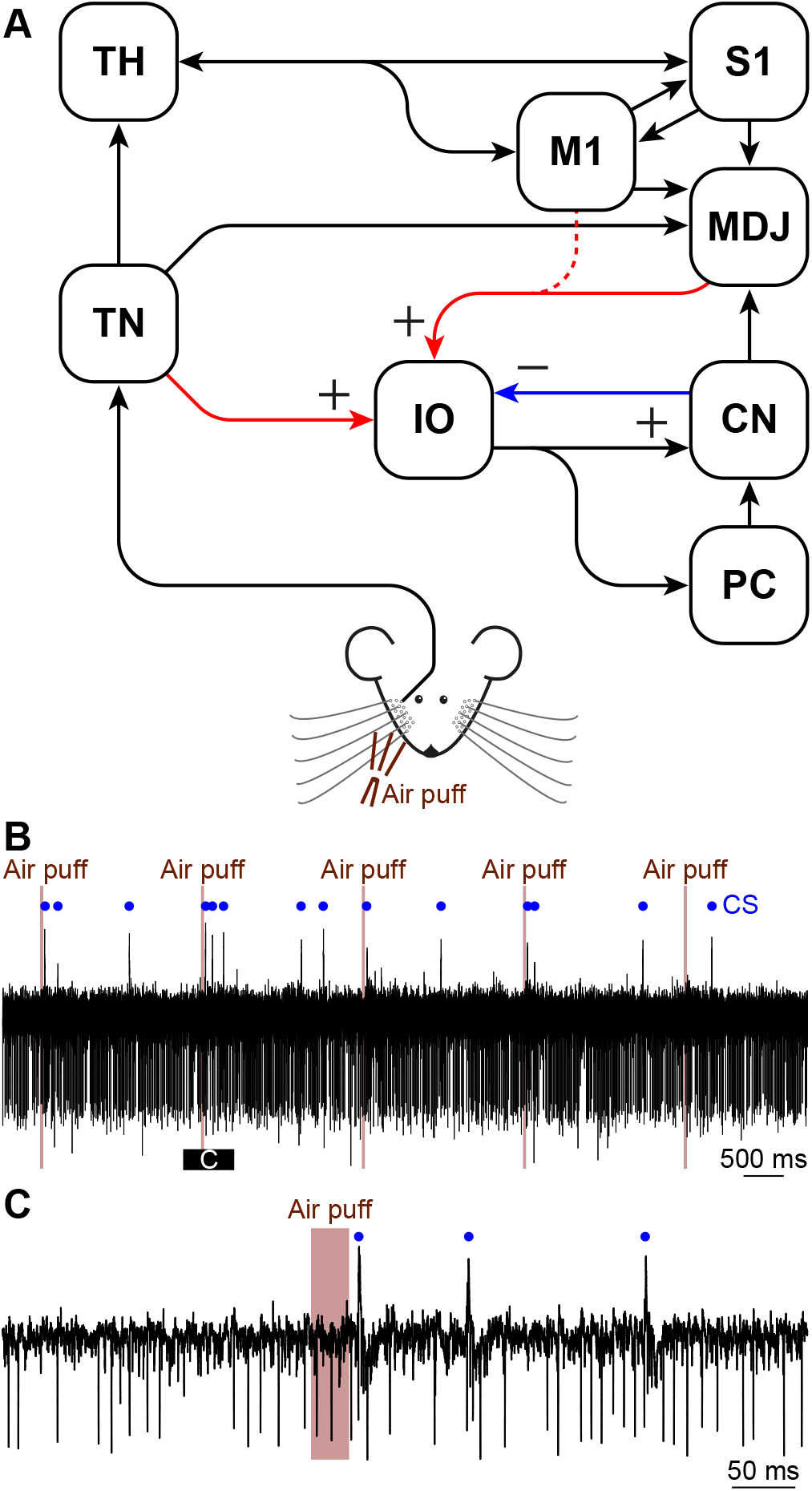
Circuit involved in the production and modulation of complex spikes. (**A**) Simplified scheme of the inputs to the inferior olive (IO). Sensory input reaches the IO directly from the brainstem and spinal cord. In our study, we used facial whisker input that is relayed via the sensory trigeminal nuclei (TN). This input is considered the “sensory input” in our modeling studies. The IO also receives continuous inputs from other brain regions, which we modeled as the ‘‘contextual input”. The contextual input consists of excitatory input from the cerebral cortex (e.g., the motor (M1) and somatosensory cortex (S1)), either directly or relayed via the nuclei of the meso-diencephalic junction (MDJ), as well as of inhibitory input from the cerebellar nuclei (CN). The output of the 10 is directed via its climbing fibers to the Purkinje cells (PCs) in the cerebellar cortex and via its climbing fiber collaterals to the CN. Sensory input also affects the contextual input indirectly, via the strong pathway from the TN via the thalamus (TH) to the cerebral cortex. (**B**) Representative trace of Purkinje cell activity showing high frequency simple spikes (as downward deflections) and occasional complex spikes (CS; marked with a blue dot). A part of the trace is enlarged in (**C**). All recordings were made in awake mice.

Here, we combine recordings in awake mice – in the presence and absence of gap junctions – with network simulations using a novel inferior olivary model to study the functional relevance of STOs in terms of resonant spikes. We are led to propose a view of inferior olivary function that is more consistent with the interplay between STOs, gap junctions and inputs to the inferior olive. Rather than acting as a strictly periodic metronome, the inferior olive appears more adequately described as a quasiperiodic ratchet, where cycles with variable short-lasting periods erase long-term phase dependencies.

## RESULTS

### Stimulation promotes oscillatory patterns in profiles of complex spikes over short periods

To study the conditions for, and consequences of, rhythmic activity of the inferior olive, we made single-unit recordings of cerebellar Purkinje cells in lobules crus 1 and crus 2 (*n* = 52 cells in 16 awake mice) in the presence and absence of short-duration (30 ms) whisker air puff stimulation (Fig. 1B-C). In the absence of stimulation, the complex spikes of 35% of the Purkinje cells (18 out of 52) showed rhythmic activity (Fig. 2A-C; S1 Fig.) with a median frequency of 8.5 Hz (inter-quartile range (IQR): 4.7-11.9 Hz). Upon sensory stimulation, 46 out of the 52 cells (88%) showed statistically significant complex spike responses. Of these, 31 (67%) had sensory-induced rhythmicity (Fig. 2D-F), which was a significantly larger proportion than during spontaneous behavior (*p* = 0.002; Fisher’s exact test). The median frequency of the oscillatory activity following stimulation was 9.1 Hz (IQR: 7.9-13.3 Hz). Hence, the preferred frequencies in the presence and absence of sensory stimulation were similar (*p* = 0.22; Wilcoxon rank sum test) (Fig. 2C, F and 3). The duration of the enhanced rhythmicity following stimulation was relatively short in that it lasted no more than 250 ms. With our stringent Z-score criterion (>3), only a single neuron showed 3 consecutive significant peaks in the peri-stimulus time histogram (PSTH) (data in Fig. 2). The minimum inter-complex spike interval (ICSI) across cells was around 50 ms, putatively representing the refractory period. We conclude that complex spikes also display rhythmicity in awake behaving mice, and that sensory stimulation can amplify these resonances in periods of a few hundred milliseconds, even though stimulation is not required for the occurrence of rhythmicity per se.

**Fig. 2.**
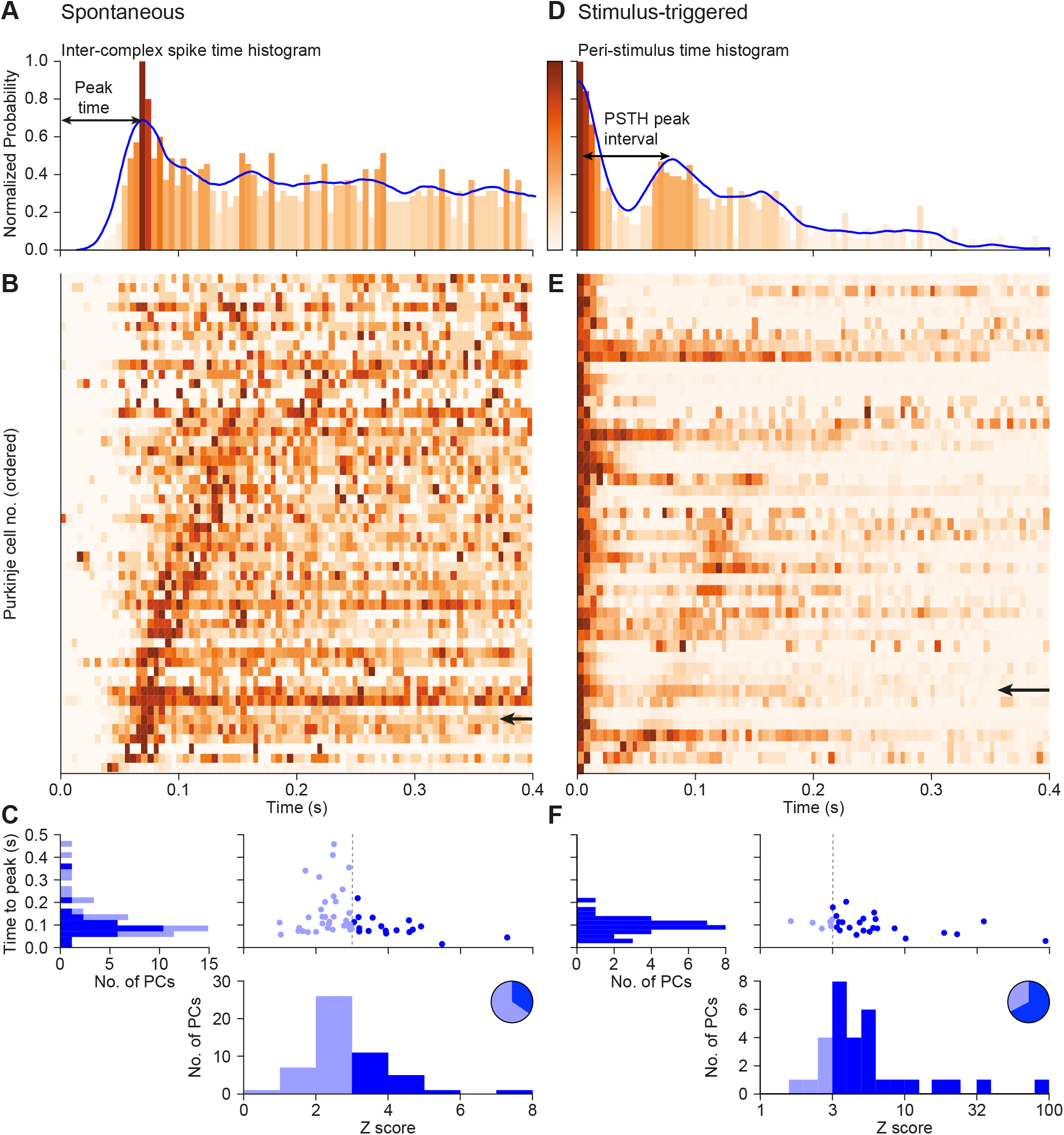
Oscillatory dynamics in complex spike firing *in vivo*. (**A**) Representative inter-complex spike-interval (ICSI) histogram of spontaneous firing of a representative Purkinje cell, together with the convolved probability density function (in blue). The colors of the bins represent the normalized oscillation strength. Note that the peak at o ms is removed to improve visibility. (**B**) Heat map showing the normalized oscillation strengths of 52 Purkinje cells, which were ordered by the time to the first side peak. The arrow in **B** indicates the cell illustrated in **A**. (**C**) The peak times of the ICSI distributions during spontaneous firing, as observed in **B**, against the Z-scores of these peaks. A Z-score <3 was taken as sign for the absence of rhythmicity (light blue fillings). The pie diagram shows the fraction of Purkinje cells with rhythmic complex spike firing (Z-score > 3; dark blue fillings). (**D**)-(**F**) The corresponding plots for the 46 (out of the 52) cells that showed a significant complex spike response following whisker air puff stimulation. Complex spike rhythmicity is displayed using peri-stimulus time histograms (PSTHs) aligned on the peak of the first response and ordered based on the latency to the second peak.

**Fig. 3.**
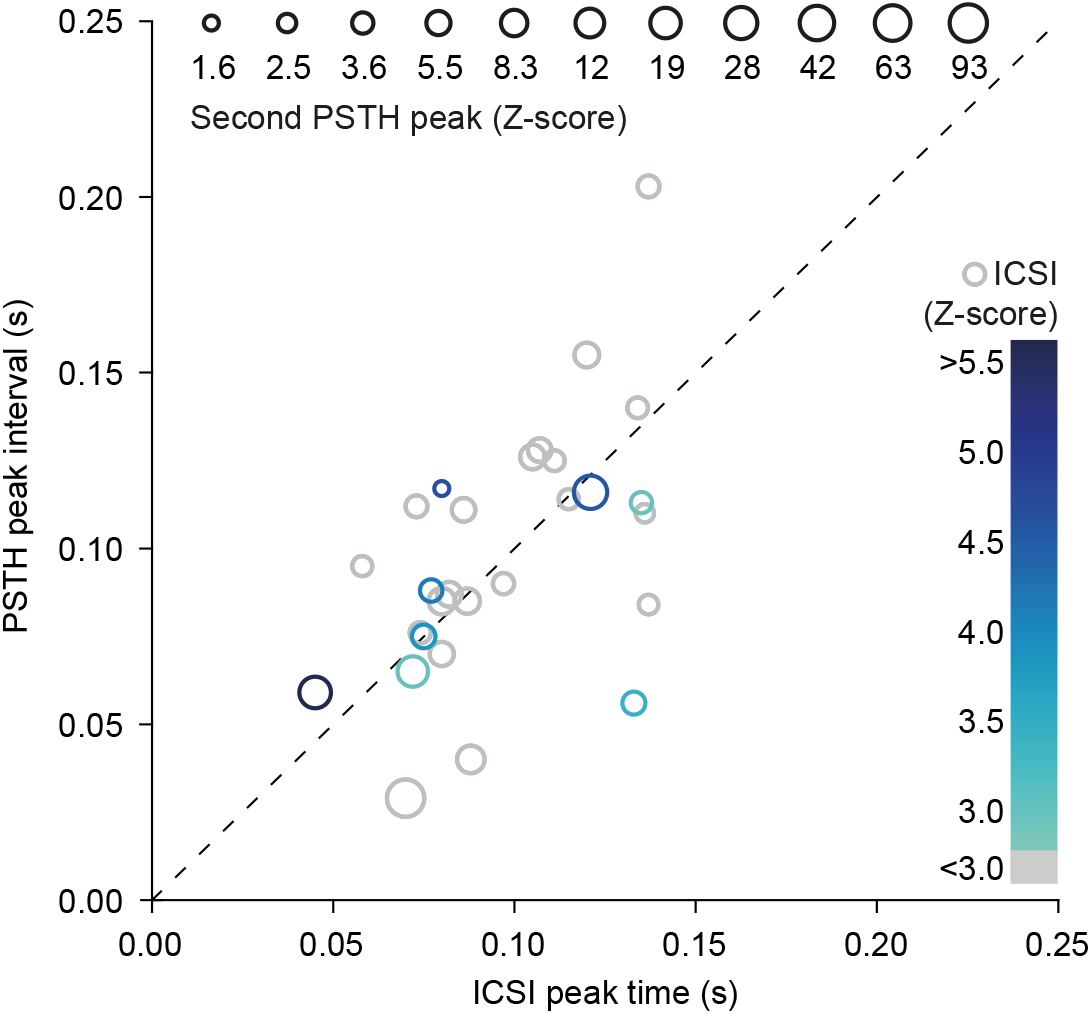
Similarity between spontaneous and sensory-induced rhythmicity. For each of the 30 oscillating Purkinje cells that were recorded in both the presence and absence of sensory stimulation, we compared the timing of the maximum of the first side peak in the inter-complex spike interval (ICSI) histogram in the absence of sensory stimulation (x-axis) with the timing of the difference between the maxima of the first and second peak in the peri-stimulus time histogram (PSTH; y-axis) following sensory stimulation. Color and size of data points indicate amplitude (in Z-score) of respectively the ICSI peak and of the second response peak. Grey scatter-points indicate the data-points falling in the low Z-score (<3) group during spontaneous firing. Note that colored circles are preferably located around the identity line.

### Rhythmic patterns of complex spikes following stimulation are transient

The pattern of rhythmic complex spike responses that was apparent for a couple of hundred milliseconds after a particular air puff stimulus repeated itself in a stable manner across the 1,000 trials (applied at 0.25 Hz) during which we recorded (Fig. 4A-B). For example, the level of rhythmicity of the first 100 trials was not significantly different from that during the last 100 trials (comparing spike counts in first PSTH peak, *p*= 0.824; χ^2^ test, or latency to first spike (*p* = 0.727, *t* test). This strongly indicates that there is – in a substantial fraction of the Purkinje cells – persistent oscillatory gating of the probability for complex spikes after a sensory stimulus resulting in time intervals (“windows of opportunity”) during which complex spikes preferentially occur (Fig. 4B-E). These windows of opportunity become even more apparent when sorting the trials on the basis of response latency: the first complex spikes with a long latency following the stimulus align with the second spikes of the short latency responses. Similarly, there are trials during which complex spikes appear only at the third cycle (Fig. 4C, seen as a steeper rise around trial no. 650). The occurrence of spikes during later cycles, not predicated on prior spikes, argues against refractory periods or rebound spiking as the sole explanations for such rhythmic firing (58) and highlight the putative existence of network-wide coherent oscillations.

**Fig. 4.**
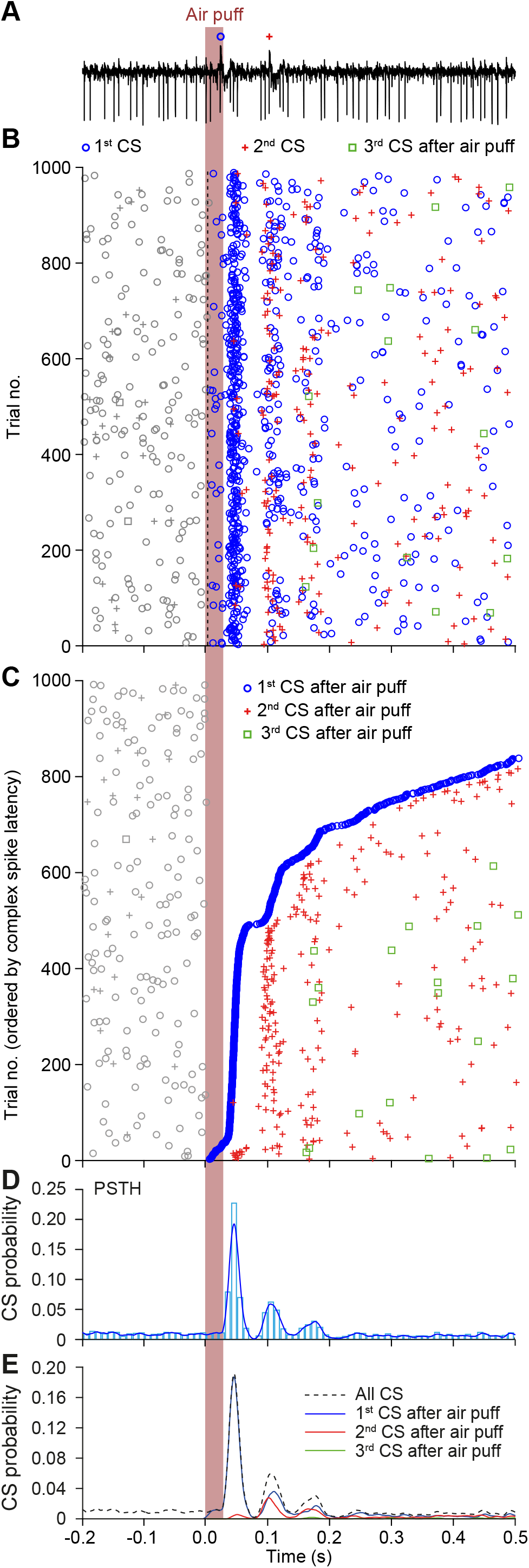
Stability of rhythmic complex spike activity in a Purkinje cell of an awake mouse. (**A**) Rhythmic complex spike (CS) responses to whisker pad air puff stimulation in a Purkinje cell of cerebellar lobules crus 1 and crus 2. Representative trial from an extracellular recording with a stimulation frequency of 0.25 Hz. In this and subsequent panels, CSs are marked with symbols representing their order of occurrence following the stimulus within each trial. (**B**) Rhythmic behavior of CSs becomes apparent in a raster plot. Note that the rhythmicity is relatively stable over the 1,000 trials. (**C**) Same as in **B** but with trials sorted by the latency to first CS. This reveals that the occurrence of CSs is largely organized in temporal windows of opportunity. Some CSs appear in later response peaks without prior firing in earlier windows of opportunity. As the CS response may appear in the early and/or in the late window, this is suggestive of a readout from an underlying oscillatory process. This phase dependence would not appear if it would be solely due to a reset followed by transient oscillation evoked exclusively in the spiking cell. If CS firing during the later peaks was predicated on an earlier phase-aligned CS, then later CS responses would have no reason to align with the second or the third peaks of the PSTH (in **D** and **E**). (**D**) PSTH of CS firing. The bin-width is 10 ms and the blue line shows the convolved histogram with a5 ms kernel. (**E**) The same PSTH as in **D**, but now the probabilities of the first, second and third CS after the stimulus shown separately, highlighting the occurrence of windows of opportunity for stimulus triggered CSs.

### Behavioral correlates of complex spikes patterns following stimulation

Since sensory stimulation of the whiskers can trigger a reflexive whisker protraction (59–61) and complex spike firing is known to correlate with the amplitude of this protraction (61), we examined the relation between periodic complex spike firing and whisker protraction. To this end, we further analyzed an existing dataset of simultaneously recorded Purkinje cells and whisker movements during 0.5 Hz air puff stimulation of the ipsilateral whisker pad. In line with our previous findings (61), trials during which a single complex spike occurred within 100 ms of whisker pad stimulation had on average a slightly, but significantly stronger protraction (from 6.1 ± 5.4° to 6.8 ± 5.3° (medians ± IQR), *n* = 35 Purkinje cells, *p* = 0.044, Wilcoxon-matched pairs test after Bonferroni correction; Fig. S2A-C). Our new analysis revealed that also the occurrence of a second complex spike had an impact on whisker protraction. This could be observed as a second period of increased protraction during trials with two complex spikes. When compared to the increase in trials with a single complex spike, this second protraction was highly significant (*p* < 0.001, Wilcoxon-matched pairs test after Bonferroni correction; Fig. S2D). The rhythmic firing pattern of complex spikes was thus reflected in the behavioral output of mice.

### STOs can explain windows of opportunity

The existence of windows of opportunity for complex spike activity is compatible with the assumption of an underlying STO, and cannot solely be explained by rebound activity without invoking circuit-wide extrinsic mechanisms. To test the implications of assuming olivary STOs, we proceeded to reproduce a detailed network with a tissue-scale computer model of the inferior olive neuropil. The model is constituted by 200 biophysically plausible model cells (40, 46, 62) embedded in a topographically arranged 3D-grid (Fig. 5A-C). It has the scale of a sheet of olivary neurons of about 10% of the murine principal olivary nucleus (cf. (63)). The model was designed to test hypotheses about the interaction between intrinsic parameters of olivary neurons, such as STOs and gap junctional coupling, and extrinsic parameters including synaptic inputs during the generation of complex spike patterns.

**Fig. 5.**
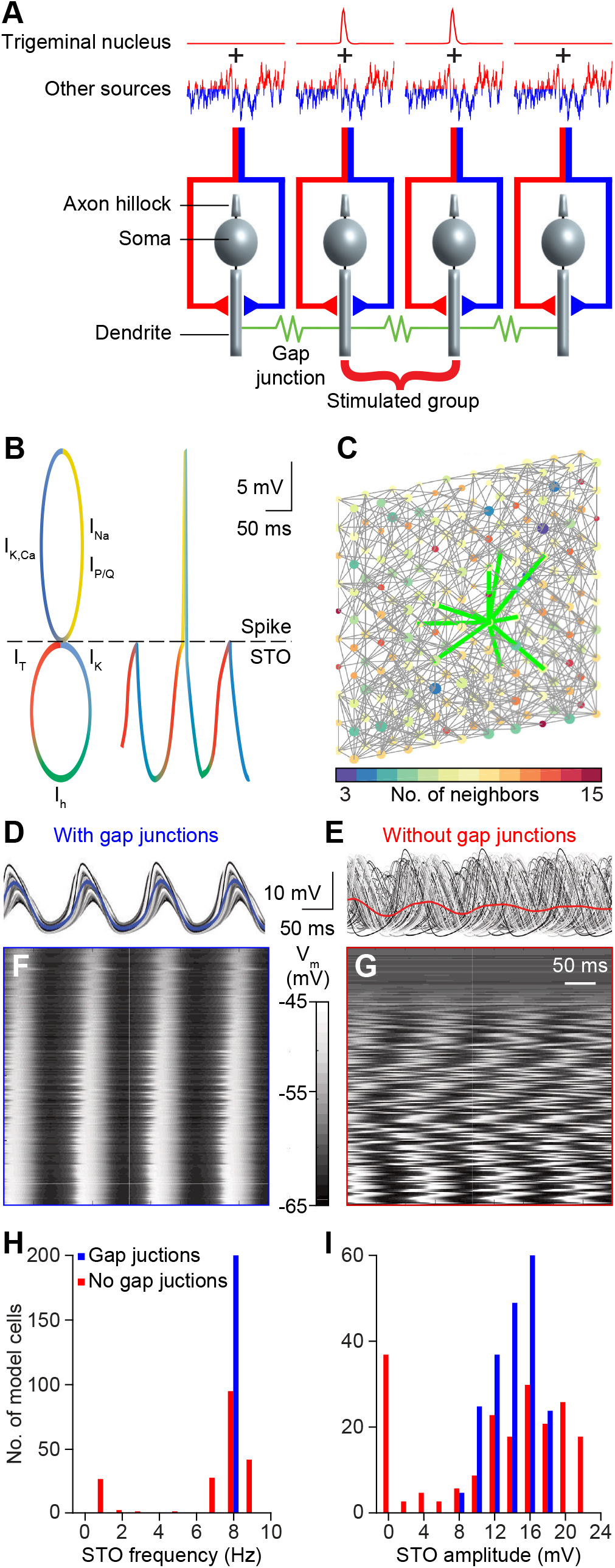
A tissue-scale network model of the inferior olive. (**A**) To study the relative impact of the different components of the inferior olive we employed a biologically realistic network consisting of 200 model neurons embedded in a 3D grid of 10×10×2. The model neurons received input from two sources: a phasic input dubbed “sensory input” and a continuous fluctuating current, emulating a “contextual input”. The “sensory input” reaches a subset of neurons synchronously and is modeled as an activation of glutamate receptor channels with a peak amplitude of 5 pA/cm^2^. The ‘‘sensory input” is complemented with a continuous input that is a mixture of excitatory and inhibitory synapses representing signals from different sources (see Fig. 1A). The ‘‘contextual input” is modeled as an Ornstein-Uhlenbeck noise process delivered to all neurons in the model network, with a 20 ms decay, representing temporally correlated input. The model neurons are interconnected by gap junctions. (**B**) The model neurons display subthreshold membrane oscillations (STOs), represented by the lower circle, and spiking, represented by the upper circle. The colors indicate which current dominates which part of the activity pattern. An l_h_ current (green), a hyperpolarization-activated cation current, underlies a rise in membrane potential; the l_τ_ current (red), mediated by low threshold T-type Ca^2+^ channels (Ca_v_3.1), further drives up the membrane potential and from here either the subthreshold oscillation continues (lower circle) or a spike is generated (upper circle), mediated through l_Na_ and l_k_ currents (yellow and blue, respectively); both after a spike or a subthreshold peak, influx of Ca^2*^ through the P/Q-type type Ca^2*^ channels (Ca_v_2.1) leads to a hyperpolarization due to Ca^2+^-dependent K^+^ channels (blue); this hyperpolarization in turn will activate the l_h_ current resuming the oscillation cycle. A resulting trace of the STO punctuated with one spike is diagrammatically displayed on the right. For a full description of currents, consult methods. (**C**) 3D connectivity scheme of the 200 neurons in the network model. Links indicate gap junctional interconnections; the color of each neuron in the model represents the number of neurons to which that cell is connected. The thick green lines indicate first order neighbors of the center cell. (**D-E**) Membrane potential of cells in the absence of input, for networks with and without gap junctional coupling. (**F-G**) Cell activity sorted by amplitude, indicating that coupling the networks brings non-oscillating cells into the synchronous oscillation. (**H-I**) Distribution of STO frequency and amplitude for coupled and uncoupled model networks.

Each neuron in the model is composed of a somatic, an axonal and a dendritic compartment, each endowed with a particular set of conductances, including a somatic low threshold Ca^2+^channel (Ca_v_3.1; T-type), a dendritic high threshold Ca^2+^channel (Ca_v_2.1; P/Q-type) and a dendritic Ca^2+^-activated K^+^ channel, chiefly regulating STO amplitudes, while a somatic HCN channel partially determines the STO period (Fig. 5B; see also Methods). The dendrites of each neuron are connected to the dendrites of, on average, eight nearby neighbors (within a radius of three nodes in the grid, representing a patch of about 400 μm × 400 μm of the murine inferior olive), simulating anisotropic and local gap junctional coupling (Fig. 5C). As the inferior olive itself, our model has boundaries which have impact on local connectivity characteristics, such as the clustering coefficient, though these did not have significant impact on the average firing rate between edge and center cells (*p* = 0.812, comparing edge and center cells, Kolmogorov-Smirnov test; S3 Fig.). The coupling coefficient between model cells varied between 0 and 10%, as reported for experimental data (45, 64, 65).

Sensory input was implemented as excitatory synaptic input, simulating the whisker signals originating from the sensory trigeminal nuclei that were synchronously delivered to a subset of model neurons. Additionally, a “contextual input” was implemented as a combination of inhibitory feedback from the cerebellar nuclei and a level setting modulating input (Fig. 1A and 5A). This contextual input is modeled after an Ohrstein-Uhlenbeck process, essentially a random exploration with a decay parameter that imposes a well-defined mean yet with controllable temporal correlations (see Methods). The amplitude of the contextual input drives the firing rate of the model neurons, which we set around 1 Hz (S4 Fig.), corresponding to what has been observed *in vivo* (28, 43, 66). Thus, our model network recapitulates at least part of the neural behavior observed *in vivo* due to biophysically plausible settings of intrinsic conductances, gap junctional coupling and synaptic inputs.

### Characteristics of STO dependent spiking in the network model

Whether a model neuron at rest displays STOs or not is largely determined by its channel conductances. Activation of somatic T-type Ca^2+^ channels can trigger dendritic Ca^2+^-dependent K^+^ channels that can induce l_h_, which in turn can again activate T-type Ca^2+^ channels, and so forth. This cyclic pattern can cause STOs that could occasionally produce spikes (Fig. 5B). In our model, the conductance parameters were randomized (within limits, see Methods) so as to obtain an approximate 1:3 ratio of oscillating to non-oscillating cells (S5 Fig.) guided by proportions observed *in vivo* (43). Sensitivity analysis with smaller ratios (down to 1:5) did not qualitatively alter the results (data and analyses scripts are available online in https://osf.io/6x5Uy/).

In the absence of contextual input, model neurons were relatively silent, but when triggered by sensory input, as occurred in our behavioral data (Figs. 2 and 4), STOs synchronized by gap junctions would occur for two or three cycles (Fig. 6A-B). Our network model confirms that gap junctional coupling can broaden the distribution of STO frequencies and that even non-oscillating cells may, when coupled, collectively act as oscillators (S6 Fig.) (67). Adding contextual input to the model network can lead to more spontaneous spiking in between two sensory stimuli. Compared to the situation in the absence of contextual input, the STOs are much less prominent and the post-spike reverberation is even shorter (Fig. 6C). Accordingly, despite the significant levels of correlation in the contextual input (10%), the periods between oscillations are more variable due to the interaction of the noisy current and the phase response properties of the network. In addition, in the presence of contextual input our model could readily reproduce the appearance of preferred time windows for spiking upon sensory stimulation as observed *in vivo* (cf. Fig. 4). This was particularly true for the model cells that directly received sensory inputs (Fig. 6D-E). Moreover, the observed rhythmicity in model cells as observed in their STO activity was in tune with that of the auto-correlogram (Fig. 6F) in that the timing of the STOs and that of the spiking were closely correlated (cf. Fig. 5B). It should be noted though that model cells adjacent to cells directly receiving sensory input showed only a minor effect of stimulation. Thus, even though the gap junction currents in the model were chosen as the ceiling physiological value for the coupling coefficient (≤10%) (45, 67), these currents alone were not enough to trigger spikes in neighboring cells.

**Fig. 6.**
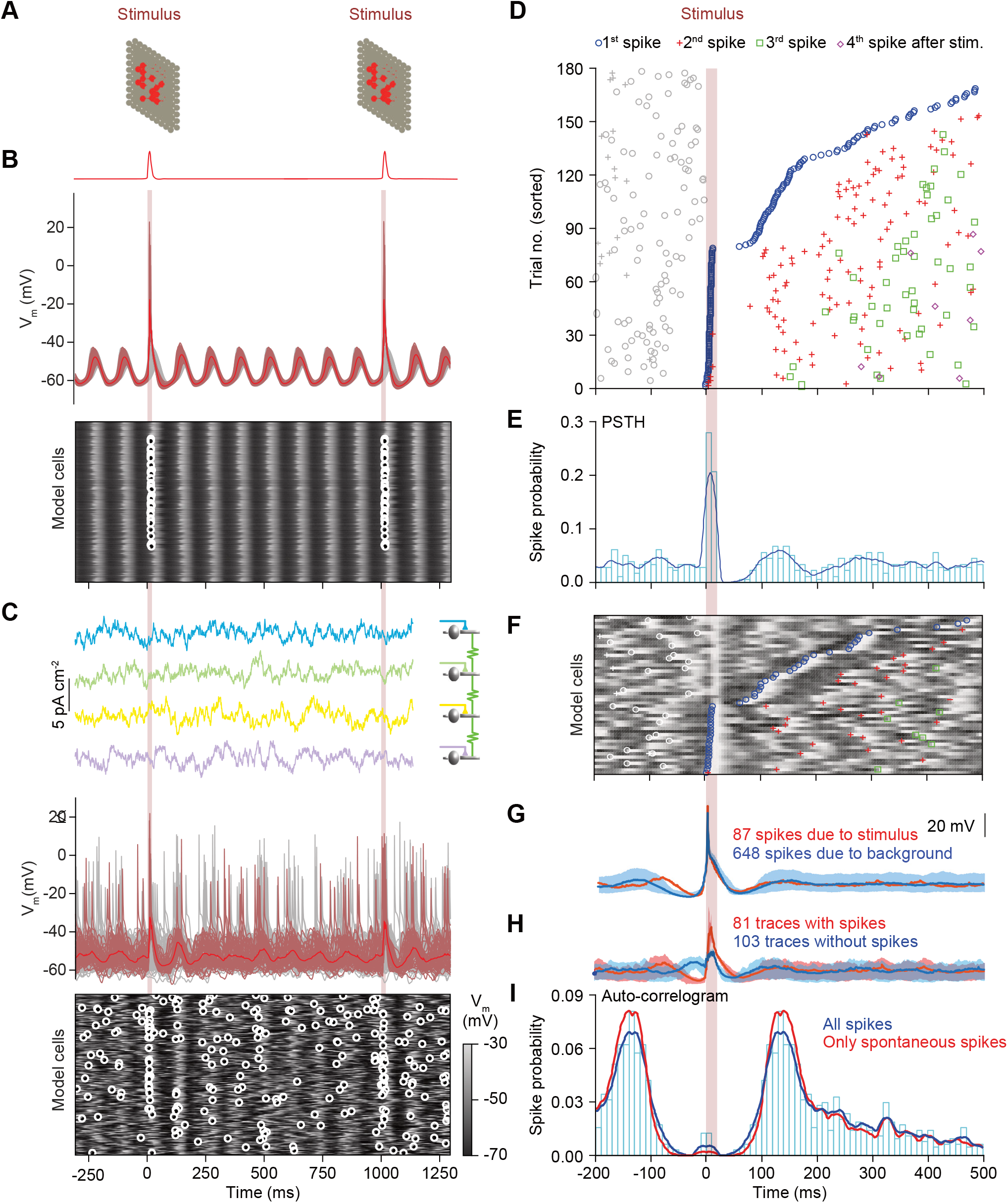
Synaptic input strongly affects the oscillatory behavior in a network model of the inferior olive. (**A**) A phasic and synchronous input mimicking sensory input (see Fig. 5A) was delivered to a subset of cells (indicated in red) of the model network. (**B**) Membrane potentials of all 200 cells in the network model in the absence of contextual input. Red trace (top) displays membrane potential average of cells receiving sensory input (see masks in **A**). Below are the membrane potentials of the model cells as a heat map. Each row exhibits the membrane potential of one model cell with the small circles indicating spikes. In the absence of contextual input, model neurons displayed long lived, synchronous and regular transient subthreshold oscillations. Note that the fact that the two stimuli fall in similar phases is coincidental. (**C**) Patterns of spiking responses changed dramatically in the presence of contextual input. Whereas in the absence of contextual input model neurons showed synchronous subthreshold oscillations and were silent except upon receiving sensory input, in the presence of contextual input the subthreshold oscillations are irregular and not strictly synchronous, while spikes occur throughout the simulation (∼1 Hz average network firing). Note that despite the largely uncorrelated (90%) contextual input, model cells did not fire homogenously as in **B**, though loose clusters of synchronously firing cells did emerge. (**D**) Rhythmic responses to stimulation are reproduced by the inferior olivary network model. A representative cell of the model network with responses to sensory input reproduces rhythmic features of the PSTH and an auto-correlogram as found *in vivo* (see Fig. 4). Raster plot of olivary spiking triggered by 182 stimuli, ordered by latency to the first spike following the stimulus. As seen in the *in vivo* data, the first spike after the stimulus can fall in multiple windows of opportunity. For this simulation, we included the contextual input. (**E**) Peri-stimulus time histogram showing the response peaks upon sensory input. (**F**) Raster plot like in **C**, but with overlaying membrane potentials. For clarity only the first 50 stimuli are displayed. Spike symbols follow conventions as in **D**. Trials with low latency were preceded by a dip in the preceding membrane potential, a phenomenon that can also be observed in **D** and **H**. (**G**) Average membrane potential of this neuron aligned by spikes that were either due to sensory input (87 spikes; red) or produced spontaneously (648 spikes; blue). The light shaded backgrounds represent the 10% and 90% percentile ranges of membrane potential. Dips in membrane potentials preceding the spikes reveal that for both groups a prior inhibition increased the probability of spiking. Importantly, except for a refractory period, we did not observe a clear oscillation pattern following either spike type, due to variability imposed by the contextual input (**H**) shows the difference in spike triggered membrane potential averages for stimulus triggered and “spontaneous” spikes. (**I**) Auto-correlograms comparing rhythmicity of spikes produced during stimulation with spikes due to background (spontaneous) activity. Periodic stimuli delivered to the network **reduced** rhythmic responses at the peak. The extreme short-latency responses (in the center of the auto-correlogram) are due to the spikelets detected in the model traces that can also be seen in **D**.

Both directly stimulated model cells and those receiving only contextual input exhibited phase preferences, seen in the spike-triggered membrane potential average as well as in the spike-triggered average of the input currents (Fig. 6G-I). Spike-triggered averages of membrane potentials for any cell showed depolarization followed by hyperpolarization. In contrast, trials in which no spike was generated showed a depolarization just before the occurrence of the input. Similarly, the average of the input showed a long-lived phase preference, not only for a hyperpolarization before the spike, but also a preference for a depolarization in the previous peak of the STO, more than 100 ms earlier. These results are in line *in vitro* experiments under dynamic clamp and noisy input (68, 69). Likewise, the model indicates that for short durations STOs can induce clear phase dependencies for spiking, which fades under the variation of period durations dependent on the trial-specific contextual input (as seen in our data).

### Phase-dependent gating in the network model

Depolarizing sensory input delivered onto a subset of the model cells can reset the STO phase in oscillatory cells and create resonant transients in others (Fig. 7A-B; see also S7 Fig. on the appearance of rebound firing). If a second stimulus is delivered during this short-lasting transient, the response probability is increased. As in most cells with resonant short-lasting dynamics, inputs delivered during different phases can cause phase advances or delays. Hyperpolarization advanced the phase between 0 − *π* and delayed the phase between *π* − 2*π*, whereas depolarization had roughly the opposite effect, in addition to phase advancements with spikes in later cycles between *π* − 2 *π* (Fig. 7C-E). Thus, there is a mutual influence of synaptic inputs and STOs on periodicity. While STOs can lead to phase-dependent gating, synaptic input can either modulate or reset the phase of the STOs, generating variable periods that range between 40-160 ms for the chosen amplitude of the contextual input (Fig. 8; S6 Fig.).

**Fig. 7.**
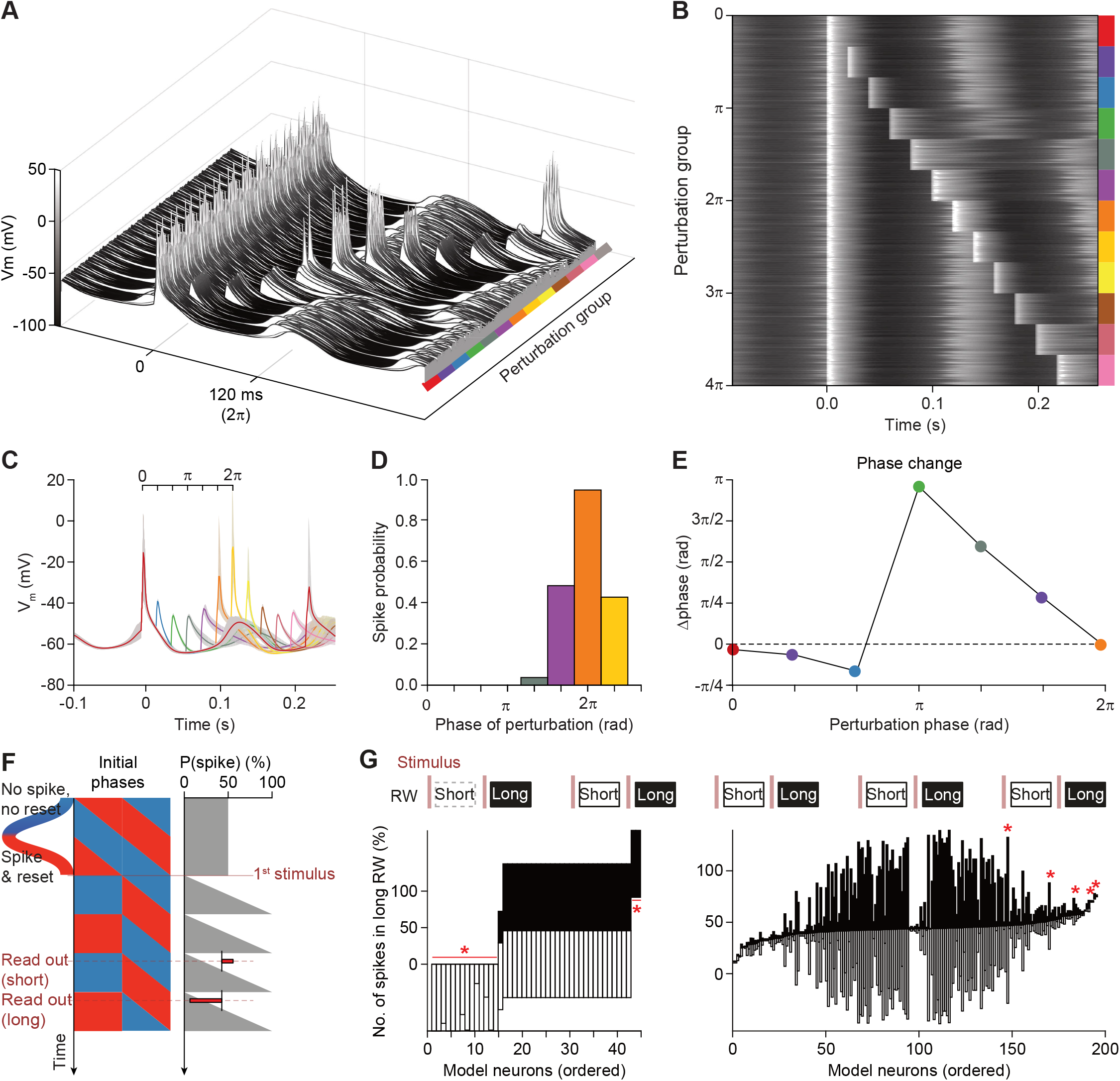
Synaptic input determines the phase-dependent gating properties of the inferior olive network model. (**A**) Phase-dependent spiking was prominent in our network model of the inferior olive in the absence of contextual input. We repeatedly stimulated the same 54 model cells with synchronous sensory input (cf. Fig. 5A). The other 146 neurons of the network model were present, but not directly stimulated. Phase-dependent gating was studied using tandem stimulation with varying inter-stimulus intervals. The membrane potential of each of these 54 model neurons is plotted in **A** as a waterfall plot and in **B** as a heat map. The simulations are grouped by the interval between the two stimuli, starting with a single stimulation (red), and continuing with two stimuli with small to large intervals. In 13 equal steps two whole cycles of the subthreshold oscillation were covered. (C) Average and inter-quartile range (shaded area) of membrane potentials of neurons in the network that were only perturbed once at 0 ms (red) or twice (other colors). The phase of the subthreshold oscillations of the “synchronized” cells in the network was used to compute the data displayed in **D** and **E**. (**D**) Spike probability plot as measured for the second perturbation at the different phases of the membrane oscillation. Spike probability was computed as the number of neurons firing divided over the total number of neurons getting input. The first perturbation given at the peak of the oscillation is demarked as the start of the oscillation cycle. Perturbations at both a half and one-and-a half cycle (π and 3π respectively) did not trigger spikes, whereas perturbations at either one or two full oscillation cycles (2π and 4π, respectively) did (see also **A** and **B**). The repetition of the ‘sinusoidal’ probability curve shows that gating is not (only) due to a refractory period, but follows the hyperpolarized phase of the oscillation (not shown). (**E**) Phase response curve showing that perturbations early in the oscillation phase did not have a large impact on the timing of the spike, but halfway the oscillation the perturbations advanced the ongoing phase considerably, with impact declining linearly to the end of the full cycle. Note that the average membrane potential at the perturbation at 0 ms only went up to −15 mV and not up to the average spike peak of approximately 10 mV, because in the population only a fraction of neurons spiked. (**F**) Gallop stimuli provide indirect evidence for an underlying oscillatory process. This idealized diagram illustrates the impact of a resetting stimulus on the future spiking probabilities. Here we assume that resetting spikes occur only during the rising phases of the STO (red) and not during the falling phases (blue). One can query for the impact on spiking probability for any given initial phase at any time after the first stimulus. Different intervals lead to biased response ratios. (**G**) In order to test the impact of the presumed phase of the inferior olivary neurons, we applied a “gallop” stimulation pattern, alternating short (250 ms) and long (400 ms) intervals (top row). “Sensory input” (vertical red shaded bars) were delivered and spikes were counted in response windows (RW) 0-180 ms post-stimulus. Response probabilities were compared between response windows (RW) in short (empty bars) and long intervals (filled bars). Many cells in the model prefer either the short or the long window (for spiking cells, see left lower panel). However, in the presence of contextual noise, this preference largely washes off, seen as approximately symmetric responses in each of the intervals. Asterisks indicate that a tiny fraction of cells still displayed significant preference for particular intervals, which could be attributed to the low number of spikes fired by these cells.

**Fig. 8.**
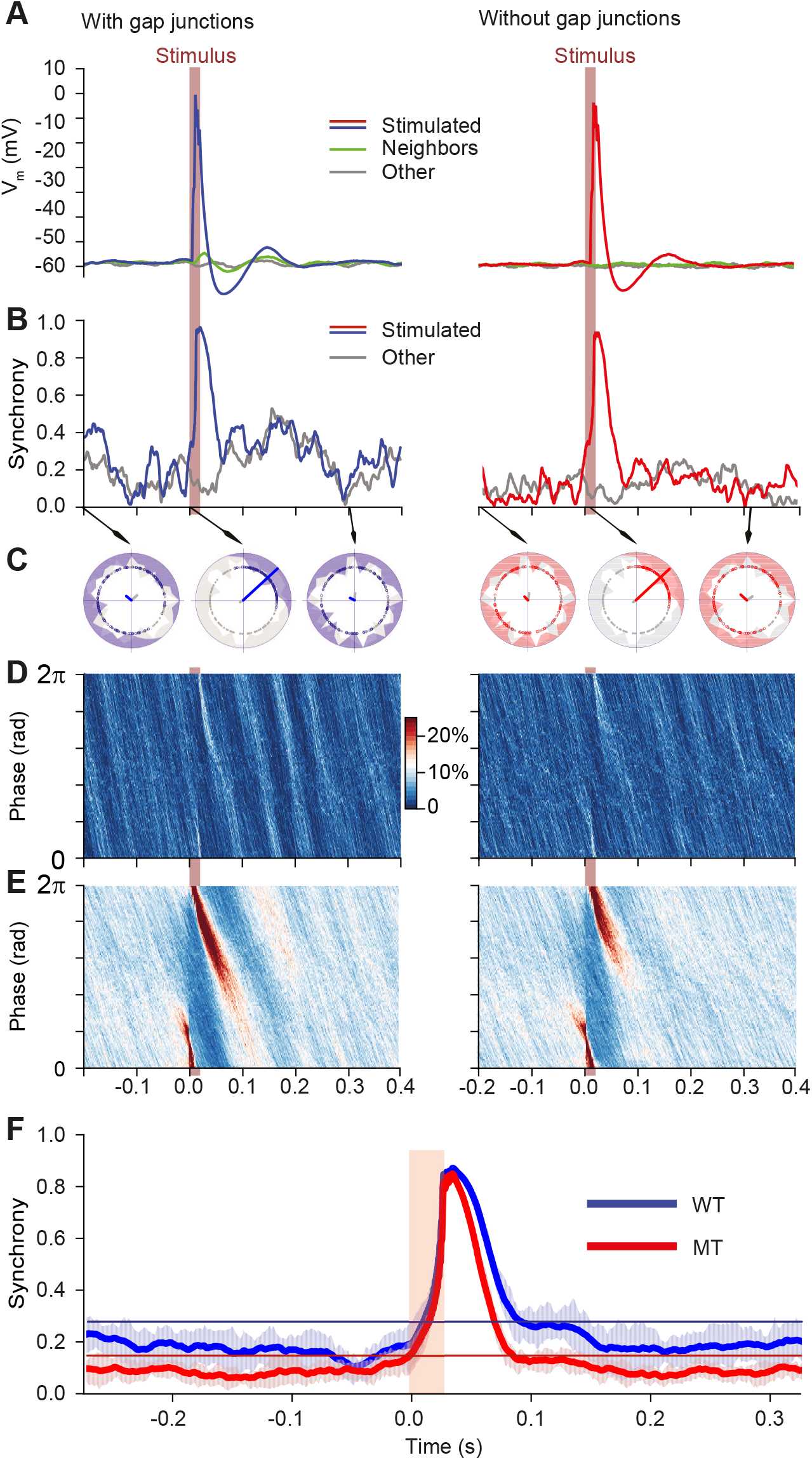
Gap junctions reinforce synchronous behavior after sensory stimulation. Impact of periodic stimulation on network phase in the presence of contextual stimuli for networks with (left column) and without gap junctions (right column). Brown bars indicate stimulation time. (**A**) Stimulus triggered membrane potential for all cells in the network (n = 100 stimuli). Stimulation causes a peak in the membrane potential of the directly stimulated cells (red line; cf. mask in Fig. 5A). The effect of the current spreading through gap junctions is visible in the cells immediately neighboring the stimulated cells (green line), whereas it is absent in the network model without gap junctions. (**B**) Increase of synchrony in the network due to a single stimulus. Synchrony is displayed as phase concentration over time (the Kuramoto parameter) for cells under the stimulation mask (colored) and other cells (gray). Stimulation generated a transient synchrony peak in both networks, but this peak was considerably higher and longer lived in cells with gap junctions, due to synchrony inducing current flows. (**C**) Polar plots indicating network phase distribution for snapshots (arrows) for stimulated (colored distribution) and non-stimulated cells (gray distribution). Phase alignment of subthreshold oscillations of the inferior olivary network model was reduced in the absence of gap junctions. The lines in the middle of the plots indicate average phases for either the stimulated cells (in color) or all cells (gray). (**D**) Phase distribution of cells over time. Color code represents the proportion of neurons occupying a certain phase at a given time (phase bin size is 2π/100, time bin is 1 ms). Dark/light bands indicate that the phase alignment propagates to most cells in the network and subsists for longer durations for the WT network. Nevertheless, contextual input overruled the phase alignment within a few hundred milliseconds (~300 ms in this case). In the absence of gap junctions, the impact of the stimulus is restricted to stimulated cells, so that the coherence induced by stimulus on the network was much smaller. (**E**) Same as in **D** but displaying averaged phase distributions for 100 stimuli. The stimulus hardly evoked an effect after 200 ms, and it did not induce any entrainment, which would have been visible as vertical bands before the stimulus. The effect of the stimulus is not stereotypical, due to dependency on network state driven by contextual input (see also Fig. g). (**F**) Average network synchrony triggered on the stimulus. The horizontal lines in the bottom plot are the 95% inclusion boundaries taken from the 5 s of spontaneous network behavior to contextual input (no periodic “sensory” pulses). The model network with gap junctions displayed after the early stimulus response curve an elevated synchrony plateau, which ended after about 200 ms, whereas the model without gap junctions hardly showed any secondary plateau. There was no preceding synchrony, indicating complete absence of entrainment to the periodic stimulus.

### Gallop stimuli provide indirect evidence for STOs

The only means of settling the question about the prevalence of STOs in awake and behaving mice would be intracellular recordings of inferior olivary neurons, which remains a daunting experimental challenge. We therefore looked for a non-invasive method that could read out, from indirect and infrequent complex spikes, the presence or absence of STOs. One such paradigm, borrowed from auditory studies (70), is the gallop stimulation paradigm that we first applied to our network model (Fig. 7F).

During a gallop paradigm, stimuli are applied in quick succession with alternating intervals in the same order as the putative frequency of the oscillation (70). Enough stimuli should be applied such that after multiple presentations the stimuli sample a uniform distribution of phases. If spikes are modulated by an oscillatory process, the presence of spikes on a short interval should be able to predict, in the next interval, the absence or presence of spikes. Indeed, if the underlying process producing spikes has oscillatory components and a relatively stable period, the probability of spikes in each interval is systematically different, which would appear as asymmetric ratios of response in the different intervals. This can be then inspected as the length of the vertical bars representing ratio of probability of spiking for long or short stimulation intervals (Fig. 7G). Thus, if the period of the STO rhythm would be regular and cause phase-dependent gating, complex spike responses following each stimulus interval are expected to show preferences for the short or long window of stimulation; these preferences were indeed observed (Fig. 7G, left). However, this scenario is idealized in that they are most prominent in the absence of noise. After adding a moderate amount of contextual input, this dependency washes off, rendering the responses in the two windows more symmetric (Fig. 7G, right).

### Model exhibits network activity that is synchronous and oscillating, but quasiperiodic

In line with the experimental *in vivo* data (e.g., Fig. 2 and 4), the olivary spike rhythmicity in the network model was steadily present over longer periods, and for a wide range of contextual input parameters (S5 Fig.). In addition, it also comprised, as in the experimental data, variations in frequency and amplitude during shorter epochs (Fig. 8 and S6). Analysis of the network parameters indicates that these latter variations in oscillatory behavior can be readily understood by their sensitivity to both the amplitude (parameter ‘sigma’) and kinetics (parameter ‘tau’, temporal decay) of the contextual input. Indeed, because of the underlying Ornstein-Uhlenbeck process, the generation of contextual input converges to a specified mean and standard deviation, but in short intervals the statistics including the average network STO frequency can drift considerably (Fig. 6C and 8). Since relatively small differences in oscillation parameters such as frequency can accumulate, they can swiftly overrule longer-term dependencies created by periodically resetting stimuli, as an analysis of phase distributions shows (Fig. 9). Thus, based upon the similar outcomes of the network model and *in vivo* experiments, we are led to propose that (1) the STOs in the inferior olive may well contribute to the continuous generation of short-lasting patterns of complex spikes in awake behaving animals, and that (2) the synaptic input to the inferior olive may modify the main parameters of these STOs. Note that in the absence of input, periodic rhythmic behavior should be the default behavior of oscillating cells. Thus, in all likelihood, even if the inferior olive oscillates endemically, sustained but variable input should induce highly contextual spike responses to variable periods and render the olivary responses quasiperiodic, rather than regularly periodic as observed in reduced preparations.

**Fig. 9.**
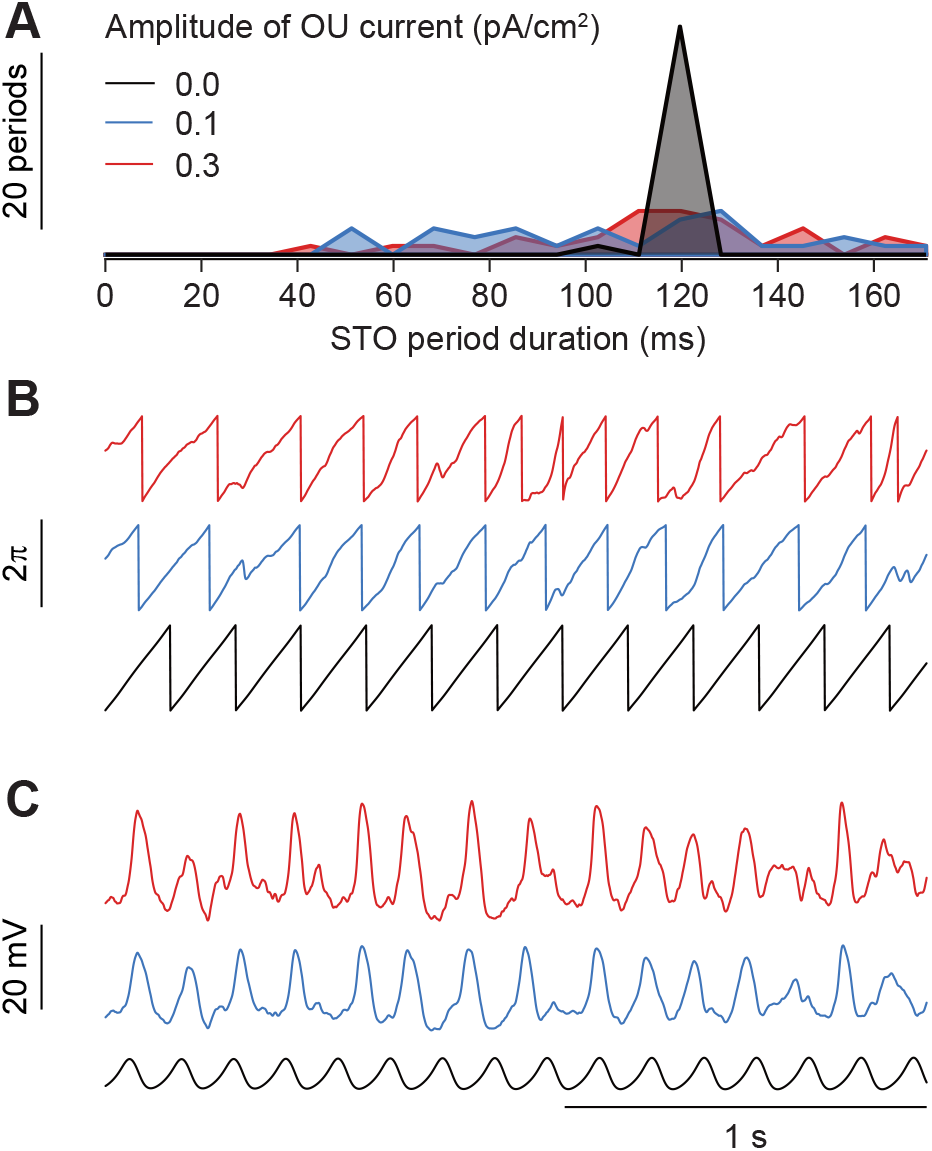
Significant variability of oscillatory periods induced by contextual input on oscillation period of a single model cell. (**A**) Distribution of inter-peak intervals for a single model cell under 5 s of contextual stimulus (OU current) of different amplitudes. To gauge period variability without introducing refractory periods, input levels were chosen for non-spiking subthreshold dynamics only. The unperturbed model cell (in black) produces regular periods, while weak levels of contextual input are sufficient to create substantial variability (red and blue). (**B**) Example traces of membrane potential and (**C**) phase for the single cell under perturbations, for the same random seed. Note that under contextual input there was substantial variation in periods, but similar STO periodicity despite different levels of input.

### Galloping stimuli show little conditional spiking in the experimental data

In line with *in vivo* whole cell recordings made under anesthesia (10, 43) our awake data support the possibility that the moment of spiking may be related to the phase of olivary STOs, especially during the period of several hundred ms following stimulation (Figs. 2 and 4). As discussed above, a gallop stimulus would expose such an oscillatory process underlying the response probabilities. However, our network model demonstrated that extrinsic input to the inferior olive could readily obscure the relation between STOs and spike probabilities (Fig. 7G). To study whether phase-dependent gating in conjunction with an underlying oscillatory process could shape complex spike response timing *in vivo* we applied both a 250 vs. 400 ms and a 250 vs. 300 ms gallop stimulation using air puffs to the whiskers. Using only trials with a CS in the previous trial to calculate the ratio of responses (‘conditional firing’) a slight bias could be observed in the 250 vs. 400 ms paradigm (Fig. 10A) and to a lesser extent in the 250 vs. 300 ms (Fig. 10B). Analysis including all trials (‘non-conditional) is included in S8 Fig. and shows no significant bias for any of the cells tested. Hence, our *in vivo* data are in line with the results from the network model and show that the timing of complex spike responses to sensory stimulation is biased but not strongly determined by STOs.

**Fig. 10.**
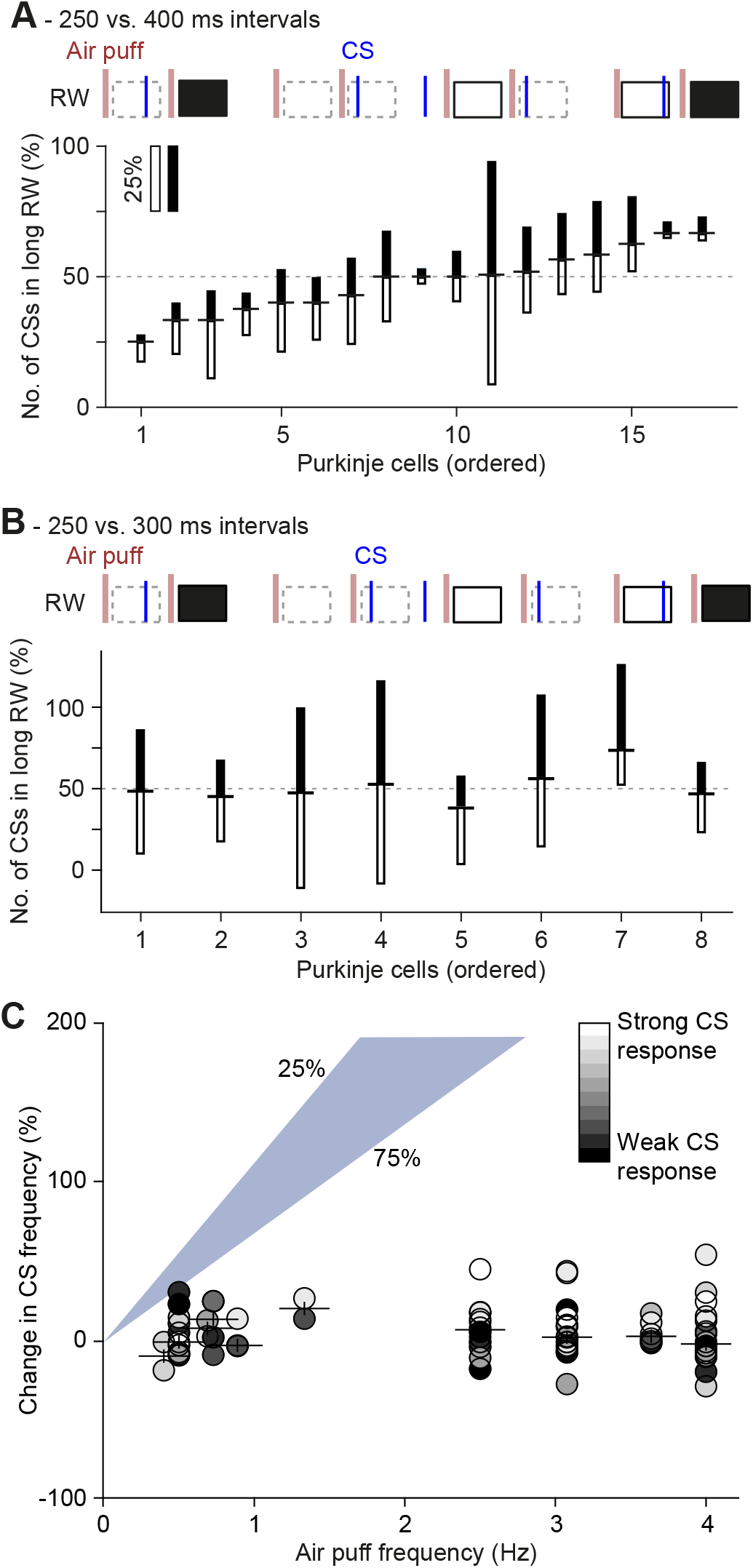
No phase-dependent spiking probabilities observed *in vivo*. (**A**) In order to test the impact of the presumed phase of the inferior olivary neurons in vivo, we applied a “gallop” stimulation pattern, alternating short (250 ms) and long (400 ms) intervals. Air puffs were delivered to the whisker pad. Complex spikes were counted in response windows (RWs) 20-200 ms post-stimulus and cells were sorted as a function of the ratio between the numbers of complex spikes in short and long intervals (indicated as horizontal dash between filled and empty bars). In this analysis, we included only the RWs that followed an RW with at least one complex spike; i.e. the RWs indicated by a dashed border were ignored. For each Purkinje cell the relative response probabilities for the long and short intervals are illustrated as the length of the filled and open bars, respectively. None of the Purkinje cells showed a significant difference in the response probability between the two intervals (all *p* > 0.05 on Fisher’s exact test). (**B**) The same for the alternation of 250 and 300 ms intervals, showing even less response bias than in **A**. (**C**) If sensory stimulation triggers complex spikes, one would expect that a higher frequency of stimulation would lead to a higher frequency of complex spikes. However, such a relation was absent. For this analysis, we included only sensory-responsive Purkinje cells of which the complex spike response exceeded a bootstrap-derived 99% probability threshold. The gray scaling of the symbols indicates the response probability. Purkinje cells displaying a higher complex spike response rate had a slightly increased firing rate upon higher stimulus frequencies, but this was far from the expected increase (based upon a linear relationship; the shaded area indicates the 25-75% range).

### High-frequency stimulation does not evoke resonant complex spike firing

Our experimental data provided evidence for phase-dependent complex spike firing during brief intervals, but gallop stimulation did not expose a strong impact of STOs on complex spike response probabilities. Therefore, we sought an alternative approach to study the impact – if any – of STOs on complex spike firing *in vivo*. We reasoned that, if a sensory stimulus triggers a complex spike response with a certain probability, higher stimulation frequency should result in a proportional increase in complex spike firing. In particular, stimulus frequencies that would be in phase with the underlying STO would be expected to show signs of resonance and result in disproportionally increased complex spike firing. However, over periods of tens of seconds the complex spike frequency was resilient to varying the stimulus frequency between 1 and 4 Hz (linear regression = −0.02; R^2^ = 0.1) (Fig. 10C) and did not show signs of resonance with any of the stimulus frequencies, as there were no frequencies at which the complex spike firing was substantially increased. Only a very high rate of sensory stimulation (10 Hz), commensurable with the average duration of windows of opportunity, could induce a mild increase in complex spike firing frequency, albeit at the cost of a highly reduced response probability (average increase: 71 ± 64% corresponding to an average increase from 1.12 Hz to 1.92 Hz; *n* = 5; *p* < 0.05; paired *t* test). This examination indicates that the average complex spike frequency is robust and stiff to modulation over longer time periods, imposing a hard limit on the frequency with which complex spikes can respond to sensory stimuli, confirming recent reports on complex spike homeostasis (71).

### STOs may contribute to complex spike firing during shorter periods

As stimulus triggered resonances were not observed at any of the stimulation frequencies, we turned to a more sensitive measure for the detection of oscillatory components in complex spike firing. We developed a statistical model that extrapolated from frequencies inferred through inter-complex spike intervals and stimulus triggered histograms (Fig. 11A-B). We reasoned that phase-dependent gating would imply that the interval between the last complex spike before and the first one after sensory stimulation aligns to the preferred frequency. In contrast, if sensory stimulation would typically evoke a phase reset, as suggested by our network model (Fig. 7), no such relation would be found. The method was applied only to Purkinje cells with highly rhythmic complex spike firing. For each of those, we calculated their preferred frequency in the absence (Fig. 11C) or presence of sensory stimulation (Fig. 11D). We used that frequency to construct statistical models representing idealized extremes of phase-dependent (oscillatory) and -independent (uniform) responses. For the oscillatory component we employed an oscillatory gating model, where the timing of the first complex spike after stimulus onset would be in-phase with the ongoing oscillation. This model was contrasted to a linear response model in which sensory stimulus could evoke a complex spike independent of the moment of the last complex spike before that stimulus, apart from a refractory period. For each Purkinje cell, we compared the distribution of the intervals between the last complex spike before and the first complex spike after stimulus onset with the predicted distributions based on the linear model, the oscillatory model and nine intermediate models, mixing linear and oscillatory components with different relative weights (Fig. 11A-E). For the two extreme models as well as for the nine intermediate models we calculated a goodness-of-fit per Purkinje cell. Overall, when using these relatively long periods (300 ms), the linear model was superior to the oscillatory model, although a contribution of the oscillatory model could often improve the goodness-of-fit (Fig. 11F-H). Despite the apparent failure of the oscillatory model to fit the data, the data did show an oscillatory profile for many of the cells (Fig. 11E). This lends support to our observations that short-lived, but reliable, oscillations are apparent in complex spike timing, although they have little impact on the timing or probability of sensory triggered CS responses.

**Fig. 11.**
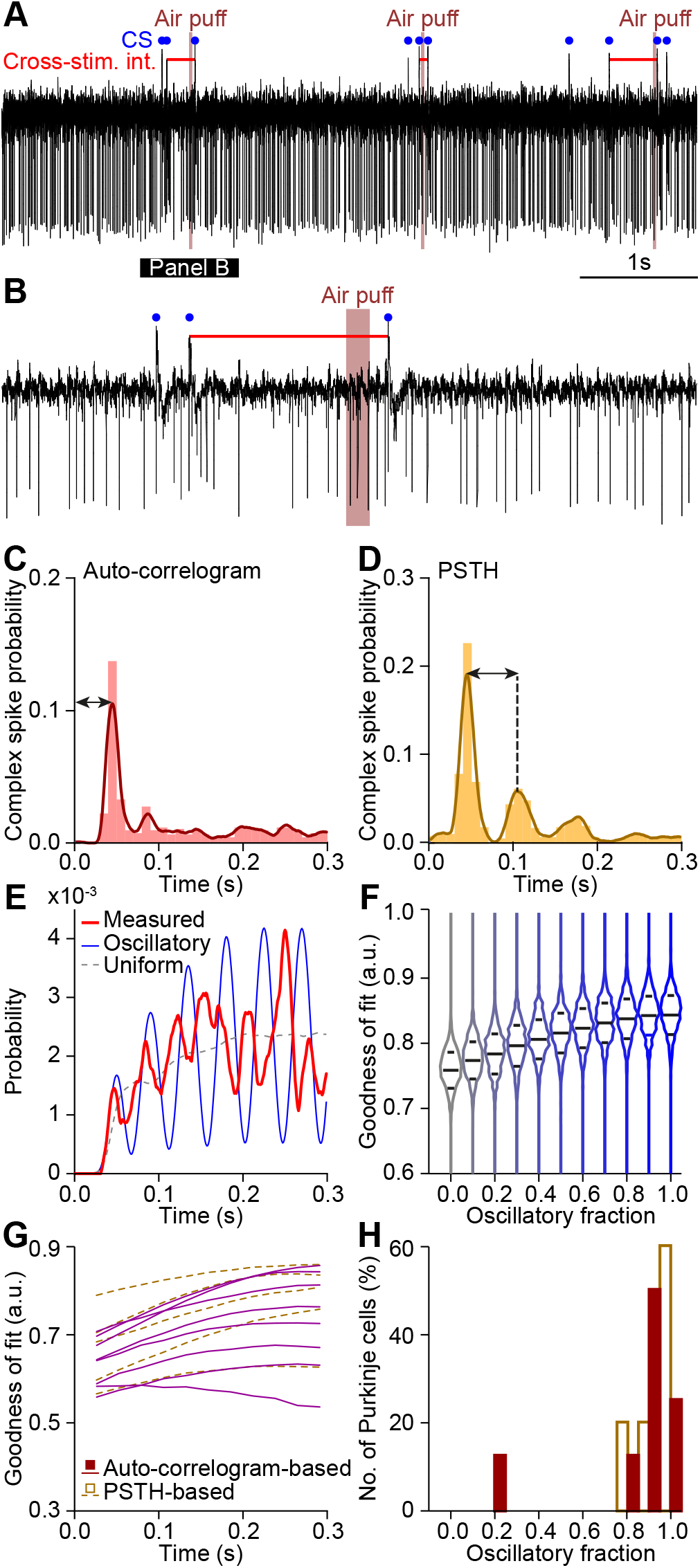
Complex spike responses to stimuli in most Purkinje cells are not conditioned on recent history. We quantified the impact of presumed subthreshold oscillations on the occurrence and timing of complex spikes by assuming a preferred firing frequency for each individual Purkinje cell according to peri-stimulus time histogram and auto-correlograms. This approach is illustrated by a representative Purkinje cell (**A-E**). A representative trace is shown in **A** and enlarged in **B**, showing the cross-stimulus interval (horizontal red line). (**C-D**) For each Purkinje cell, the preferred frequency was derived from the auto-correlogram (**C**) and the peri-stimulus time histogram (PSTH; **D**; cf. Fig. 2). (**E**) Intervals between the last complex spike before and the first complex spike after stimulus for each trial yielded a model of the preferred response windows (red line). The observed probability density function was compared with a probability density function based on a uniform complex spike distribution (dotted line), an oscillatory complex spike distribution (blue line), and 9 intermediate mixed models (see Methods). (F) The distributions of the goodness-of-fit for each of the 11 models showed a clear bias towards the uniform model, casting doubt on the impact of subthreshold oscillations on sensory-induced complex spike firing. The middle line indicates the average of all runs, while the upper and lower lines indicate 75 and 25% quartiles, respectively. (**G**) Distributions of the goodness-of-fit of all Purkinje cells that showed clear rhythmicity (see Methods). (**H**) Histograms of the best mix model shown in **G** indicate that the impact of the subthreshold oscillations on sensory complex spike responses is present, though small.

### Gap junctions facilitate frequency modulation of rhythmic complex spike firing

Apart from the STOs, extensive gap junctional coupling between dendrites is a second defining feature of the cyto-architecture of the inferior olive (31, 33, 72). Absence of these gap junctions leads to relatively mild, but present behavioral deficits in reflex-like behavior and learning thereof (10, 73). We analyzed the inter-complex spike interval times in Purkinje cells of mutant mice that lack the Gjd2 (Cx36) protein and are hence unable to form functional gap junctions in their inferior olive (69). In line with the predictions made by our network model (Fig. 5H and S6), the absence of gap junctions did not quench rhythmic complex spike firing during spontaneous activity (Fig. 12A). In fact, the fraction of Purkinje cells showing significant rhythmicity in the Gjd2 KO mice was larger than that in the wild-type littermates (Gjd2 KO: 38 out of 65 Purkinje cells (58%) vs. WT: 15 out of 46 Purkinje cells (33%); *p* = 0.0118; Fisher’s exact test), with their average rhythmicity being significantly stronger (*p* = 0.003; Kolmogorov-Smirnov test), measured by Z-scores of side peaks (Fig. 12B). Indeed, the variation in oscillatory frequencies across Purkinje cells of the mutants was significantly less than that in their wild-type littermates in that the latency to peak times per Purkinje cell were less variable (*p* = 0.0431; Mann-Whitney test; Fig. 12C). This latter finding is at first sight contradictory to our findings in the network model, where we show that gap junctions promote more uniform firing rates through increased synchrony between neurons (Fig. 5H). These simulations were run in the absence of synaptic input, though. Addition of contextual input also creates more variability in the wild type cells (S6B Fig.). As the lack of gap junctions increases cell excitability (10, 69), it is likely that synaptic input has a larger impact in the absence of gap junctions, leaving less room for inter-cell heterogeneity. Overall, removal of gap junctions affected the temporal and spatial dynamics by increasing the stereotypical rhythmicity of complex spike firing.

**Fig. 12.**
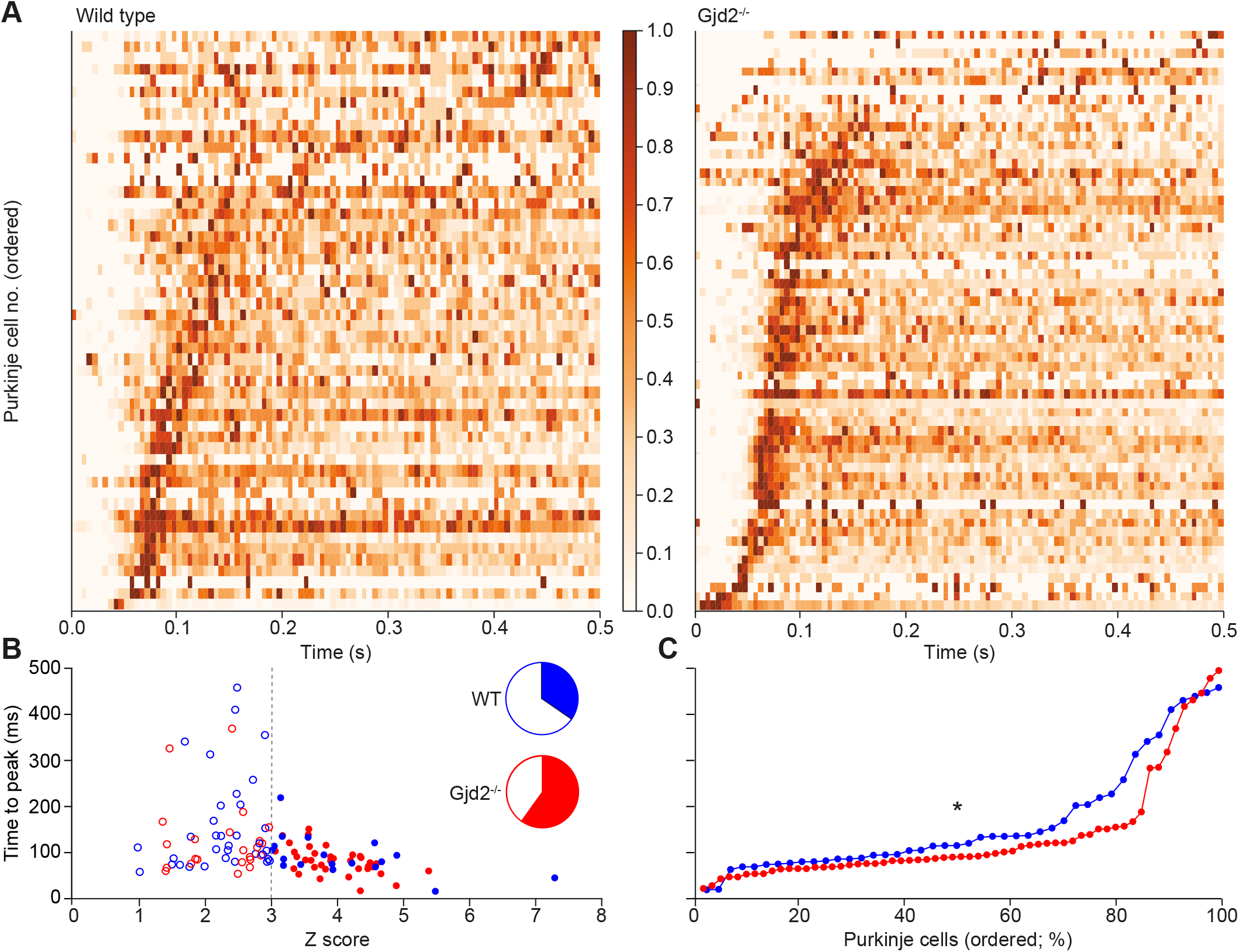
Oscillatory dynamics in spontaneous firing in vivo in the presence and absence of gap junctions. (**A**) Color-coded inter-complex spike-interval (ICSI) histogram of spontaneous firing of 60 Gjd2^−/−^ and 52 wild-type Purkinje cells. (**B**) The Gdj2^−/−^ Purkinje cell distribution shows higher Z-scores than the wild-type population, indicating a stronger rhythmicity during spontaneous firing in the absence of gap junctions (*p* = 0.003; Kolmogorov-Smirnov test). A Z-score >3 was taken as sign for the presence of rhythmicity - which occurred more often in the Gjd2 KO than in the VZT Purkinje cells (coloredfraction in the pie diagrams; *p* = 0.005; Fisher’s exact test). In the absence of gap junctions, rhythmic firing was more stereotypical, as illustrated by less variation in the time to peak (**C**; *p* = 0.0431; U = 1030.0; Mann-Whitney test). Note that the left panel of **A** is a copy of Fig. 2B and presented here to facilitate comparison.

### Temporal relations of complex spikes of different Purkinje cells

We made paired recordings of Purkinje cells in awake mice to study the temporal relations of their complex spikes during spontaneous activity. The cell pairs were recorded with two electrodes randomly placed in a grid of 8 × 4, with 300 μm between electrode centers. Cell pairs showed coherent activity in that they could show a central peak and/or a side peak in their cross-correlogram (Fig. 13A-C). The side peaks could appear at different latencies, similar to the range observed in auto-correlograms of single Purkinje cells (cf. Fig. 2B). Moreover, Purkinje cell pairs that did not produce signs of synchronous spiking in the center peak could still produce an “echo” in the side peak after 50-150 ms. Counter-intuitively, cross-correlograms of Purkinje cell pairs of the wild type mice showed less often a significant center peak than those of Gjd2 KOs (WT: 51 out of 96 pairs (53%; *N*= 4 mice); Gjd2 KO: 44 out of 61 pairs (72%; *N* = 7 mice); *p* = 0.0305; Fisher’s exact test). In line with the more stereotypic firing observed in single cells in the absence of gap junctions (Fig. 12), the strength of the center peak was on average enhanced in the mutants (*Z* scores of significant center peaks (median ± IQR): WT: 3.47 ± 1.82; Gjd2 KO: 5.75 ± 5.58; *p* = 0.0002; Mann-Whitney test) (Fig. 13D-F). Instead, the side peak of Gjd2 KO Purkinje cell pairs was not stronger than that of WTs (*Z* scores of significant side peaks (median ± IQR): WT: 3.01 ± 0.89; Gjd2 KO: 3.04 ± 1.52; *p* = 0.194; Mann-Whitney test), leading to a lower ratio between center and side peak (mean ± SEM: WT: 90.70 ± 5.17%; Gjd2 KO: 72.86 ± 6.52%; *p* = 0.036; *t* = 2.143; df = 67; *t* test).) (Fig. 13E-F). Interestingly, the occurrence of side peaks in Purkinje cell pairs was unidirectional in approximately half the cell pairs (WT: 47 out of 82 pairs with at least one side peak (57%); Gjd2 KO: 25 out of 47 pairs (53%); *p* = 0.714; Fisher’s exact test), which means that one of the neurons of a pair was leading the other, but not *vice versa*. As this was consistent in the Gjd2 KO as well as the WT Purkinje cells, these data could reflect traveling waves across the inferior olive, which, however, must have extrinsic sources (44, 74). Thus, the paired recordings are compatible with the findings highlighted above in that the presence of coupling can affect the coherence of STOs for short periods up to a few hundred milliseconds, while leaving the window for later correlated events open.

**Fig. 13.**
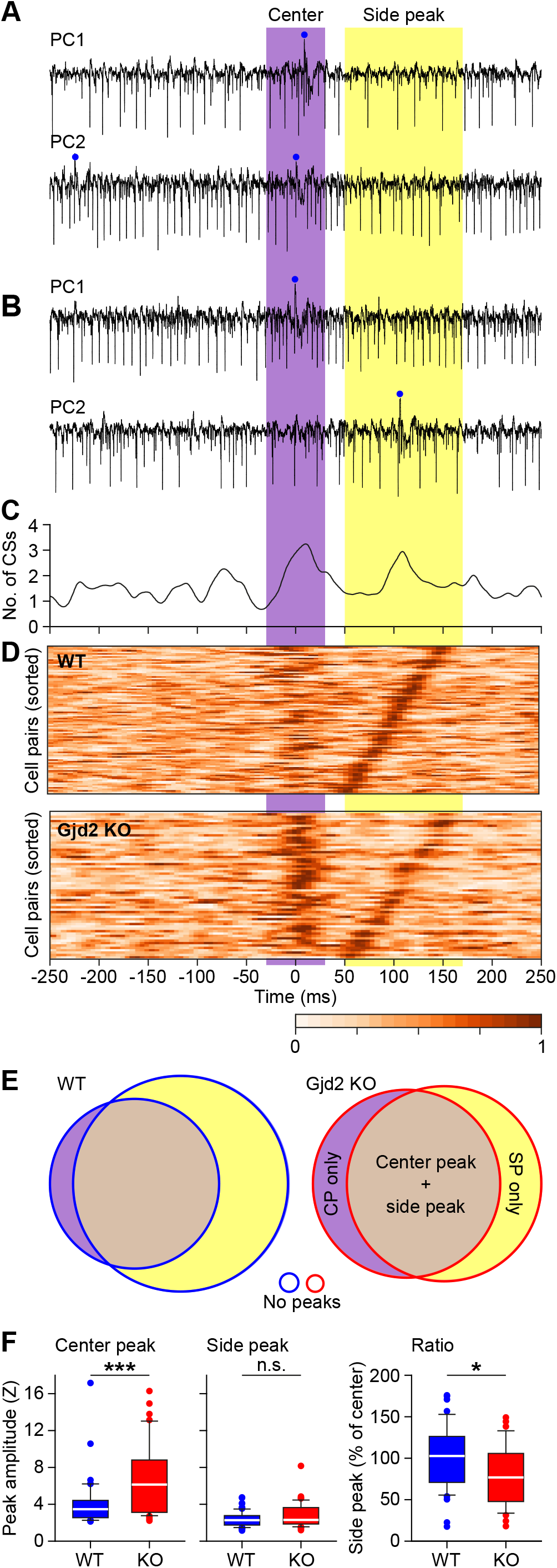
Coherence of complex spike firing of Purkinje cells over time. (**A**) Two Purkinje cells recorded simultaneously in a wild type (WT) mouse during spontaneous activity, showing an epoch with near-synchronous complex spike firing with both spikes in the center-time interval. (**B**) Another epoch of the same cell pair, but now the second Purkinje cell fired a complex spike at the next cycle, leading to a double-peaked cross-correlogram for this cell pair, seen in **C**. (**D**) Heat map representations of cross-correlograms between simultaneously recorded Purkinje cells in WT and Gjd2 KO mice. The latency of the strongest side peak was used to order the cell pairs. Correlation direction was selected so that the strongest side peaks were always positioned on the right side. Note that while the center peak is stronger in Gjd2 KO, the side peak is more prominent in WT pairs. This side peak can be regarded as a “network echo”. (**E**) Venn diagrams showing the relative occurrence of center and side peaks in the presence and absence of gap junctions. (**F**) In the absence of gap junctions, the center peak, but not the side peak, was stronger (left and middle panel). Likewise, the normalized ratio of the side peak vs that of the center peak was lower in the Gjd2 KO mice (right panel). * and *** indicate *p* < 0.05 and *p* < 0.001, respectively (*t* test).

## DISCUSSION

Given the major impact of complex spikes on Purkinje cell processing and motor behavior (10, 11, 25, 75, 76), resolving the mechanisms underlying their timing is critical to understand the role of the olivo-cerebellar system in motor coordination and learning (20, 22, 24, 77). Synchronized subthreshold oscillations (STOs) of olivary neurons have been suggested to contribute to the formation of temporal patterning of complex spike firing, but most of the evidence for STOs has been obtained in decerebrate or anesthetized animals *in vivo* or, even more indirectly, *in vitro* (10, 41–43, 45, 47, 78), but see (79). Our experiments and model were designed to inquire the existence of an *in vivo* STO, and on its ability to display phase-dependent responses in the awake brain. We evaluated rhythmic complex spike firing in behaving animals responding to peripheral stimuli and investigated their match with simulations of a tissue-scale model of the inferior olivary network. The model and data would be consistent with STOs whose period can be readily adjusted upon synaptic fluctuations from other brain regions, an effect that is consistent with the known response properties of the inferior olive (46, 67).

### Complex spike patterns are explained by cell physiology and network-wide properties

During spontaneous activity, Purkinje cells generally fire a complex spike roughly once a second, but this frequency can be increased to about 10 Hz by systemically applying drugs, like harmaline, which directly affect conductances mediating STOs in the inferior olive (41, 80). Since these drugs also induce tremorgenic movements beating at similar frequencies, it has been proposed that the inferior olive may serve as a temporal basis for motor coordination (11, 81). One concrete proposal postulates that the oscillatory properties of the olivary neurons could mirror limb resonant properties and act as an inverse controller, for example by minimizing undesirable limb oscillations (82).

In line with previous recordings (22–24, 28, 61, 71, 75, 77, 83), the current data indicate that only a small fraction of Purkinje neurons respond to sensory stimulation with an increased complex spike response probability larger than 50%. This probability falls substantially with increasing frequency of stimulation, as the overall spike frequency only marginally increases to high frequency stimulation. Even after applying different temporal patterns of sensory stimulation for longer epochs, we observed no substantial deviation from the stereotypic 1 Hz firing rate. Furthermore, our attempts to harness resonant firing by using specially designed paradigms did not evince resonant frequencies or conditional dependencies across trials. In our study, complex spikes remain as unpredictable as ever. Thus, regulatory mechanisms keep the complex spike rate relatively stable over longer time periods (71), and prevents resonance, irrespective of an enduring powerful sensory stimulus in a variety of frequencies. Save few exceptions, the presence of a complex spike in an interval is compensated by the absence in another. Thus, it looks as if the complex spikes rearrange themselves in time in order to keep close to the 1 Hz frequency. It remains to be shown to what extent the homeostatic mechanisms involved are intrinsic (cell-dependent) and/or extrinsic (network-dependent). A possible candidate for setting the overall level of excitability through intrinsic mechanisms is given by Ca^2+^-activated l^−^ channels, which are prominently expressed in olivary neurons along with Ca^2+^-dependent BK and SK K^+^ channels (84, 85). In addition, the olivo-cerebellar module itself could partly impose this regulation (85–87). Indeed, the long-term dynamics within the closed olivo-cortico-nuclear loop may well exert homeostatic control, given that increases in complex spikes lead to enhanced inhibition of the olive via the cerebellar nuclei (20). The impact of such a network mechanism may even be more prominent when changes in synchrony are taken into account. We propose that a closed-loop experiment conducted while imaging from a wide field, producing stimulation as a function of the degree of complex spike synchrony, could show conditional complex spike probabilities. Increasing our capability of predicting complex spikes is instrumental to elucidate the homeostatic control of inferior olivary firing.

### Inferior olivary STOs

The existence of temporal windows of opportunity for complex spike responses following sensory stimulation highlighted a potential impact of STOs on conditional complex spike gating (10, 41, 43, 88). Indeed, autocorrelogram peaks correlated well with interspike intervals following stimulation, arguing for an underlying rhythm. Complex spikes could appear in a particular window even when they were not preceded by a complex spike in a previous window during a single trial, arguing against a prominent role of refractory periods in creating rhythmic complex spike responses. Comparing actual firing patterns with statistical models mixing linear or oscillatory interval distributions indicated a potential impact of oscillations. The mild impact of the oscillatory component on explaining the data may in part depend on the assumption that cells have a well-defined frequency. In other words, a variable rebound time could offset the phase response by a couple of milliseconds, reducing the contribution of the oscillatory model, though phase preferences due to prior spikes may still occur (i.e., Fig. 11E). Our biophysical model suggests that fluctuating inputs, such as those mediating inhibition from the cerebellar nuclei or those relaying depolarizing modulation from the raphe nuclei (89, 90), induce variations in the oscillation period on a cycle-by-cycle basis (Figs. 8 and 9). As these contextual inputs are absent or suppressed in decerebrate or anesthetized preparations, as well as *in vitro*, they may also explain why many earlier studies systematically encountered cells with well-defined STO frequencies (10, 41, 43, 45, 47, 78, 91, 92). In the network model, in which we mimicked the contextual input as an Ornstein-Uhlenbeck process (93), the results agree well with the experimental observations in terms of synchronous firing, phase shifts, cross-correlogram peaks and side peaks, as well as overall firing frequency. Indeed, the absence of resonant responses over longer time windows and the inconsistency of individual olivary cells to fire on every trial or cycle indicate that the STOs are not regularly periodic, but rather quasiperiodic, while still being synchronous.

### Can rebound firing explain rhythmic complex spike responses?

Even though several lines of evidence suggest a role for STOs (see above), we did not observe an unequivocal, significant conditional dependence of complex spikes in the gallop paradigm, as expected by a noiseless model. How can a system with rhythmic responses at least partially fail to be phase modulated by such periodic stimuli? An attractive alternative explanation for rhythmicity might be the occurrence of Ca_v_2.1 high-threshold dependent rebound spikes (S7 Fig.) (12, 62). If impulse-like input to the olive can evoke a spike, and if this spike produces a rebound spike some tens of milliseconds later, this could explain the alignment between the PSTHs and cross-correlograms. However, this argument cannot explain stimulus triggered spikes at the second or third window of opportunity, without an earlier spike as observed in Fig. 4. As the occurrence of the rebound spike is predicated on a prior spike, a spike in the second or third window without a prior spike cannot be explained by the rebound spiking phenomenon, at least not within the same cell. In other words, the spikes happening exclusively in the second (or third) window of opportunity cannot be the result of a previous spike in the same cell, unless there is a shared rhythm in the network. It is also conceivable that strong hyperpolarization that is synchronized with the CS rhythm could promote reverberating firing, but this is an extrinsic mechanism, discussed below. As they stand, our findings do not support the idea that the post-spike hyperpolarization is a prerequisite for the complex spike pattern observed. Multiple windows of opportunity could, according to our model, be enhanced by transient oscillations induced by resets relayed by gap junctions to the local olivary circuit.

### The role of gap junctions in complex spike timing

Apart from the almost complete absence of interneurons, the presence of STOs and the exclusive projection to the cerebellum, the abundance of dendro-dendritic gap junctions is another defining feature of the inferior olive. The absence of these gap junctions does not lead to gross motor deficits, but prevents proper acquisition and execution of more challenging tasks (10, 16, 73), which is in line with the relatively minor impact found on complex spike activity in Gjd2 KO mice. At first sight, the effects of deleting gap junctions seem counterintuitive. Synchronous and rhythmic patterns are exacerbated, rather than diminished by the loss of gap junctions. However, the side peak of the auto-correlogram is significantly squashed, indicating that the gap junctions have a role in synchronizing the upcoming oscillation. Gap junctions do not only facilitate synchronization of coupled neurons, they also lower their excitability by increasing the membrane resistance (69). This results in less direct coupling, observed as reduced synchrony of direct neighbors (16, 94), and increased responsiveness to synaptic input. This leads to more long-range coherence and as a consequence gap junction networks may act as a “noise filter”, promoting short-range quorum-voting on phase (a term coined by Winfree (95)). This effect is visible in our model as spikes are most likely to occur when excitation follows inhibition (Fig 6H). This is in line with the finding that complex spikes of nearby Purkinje cells have strong preference to fire together even in the absence of sensory input (96, 97). This concept also agrees with the possibility that coupled olivary neurons may control movements by dampening the dynamics of the muscles involved at an appropriate level (82, 98), as both the resonances and movement oscillations increase shortly after sensory stimulation in Gjd2 KO mice (16).

### Reverberating loops

Network resonances are a pervasive feature of brain circuits and they can be induced by subthreshold oscillations of particular cell types (99, 100). In addition to the autochthonous dynamics of the inferior olive, reverberating loops through the circuit could help explain some features of complex spike firing, including the occurrence of complex spike doublets and side peaks in cross-correlograms. Such phenomena could be explained by “network echoes”, where complex spikes in one cycle would induce complex spikes in the next cycle (86, 101, 102). The most obvious candidate loop to produce is that via the cerebellum and the nuclei of the meso-diencephalic junction (55, 103). The output of the inferior olive is mainly directed via exceptionally strong synapses to the Purkinje cells (104). These Purkinje cells in turn inhibit neurons of the cerebellar nuclei that can show rebound firing after a period of inhibition (87, 105). This rebound activity can excite the inferior olive again via a disynaptic connection via the nuclei of the meso-diencephalic junction. While an isolated complex spike is unlikely to evoke such a rebound activity, a larger group of Purkinje cells could be successful in doing so (20, 105, 106). The travel time for this loop (around 50-100 ms) has been indirectly assessed in the awake preparation (8, 10, 26), and corresponds to the latency of the rebound firing in the cerebellar nuclei under anesthesia (86, 87, 105, 107). This implies that the travel times for the entire loop would be in the same order as found for the preferred frequencies of complex spike firing. Other, more elaborate loops involving for instance the forebrain may also exist (108) and could play an additional role in shaping complex spike patterns.

A putative impact of reverberating loops on rebound activity could be a network phenomenon, as the impact of an isolated complex spike may not be sufficient to trigger this loop. This is in line with the reduced “echo” in the cross-correlograms of the Gjd2 KO mice and enhanced doublets following lesions of the nucleo-olivary tract as occurs in olivary hypertrophy (109). Taken together, rebound spiking, STOs and reverberating loops all seem to promote in a cooperative manner complex spike rhythmicity at a time scale of about 200 ms. Through modeling, we found that not only the state of the inferior olivary oscillations determines which inputs are transmitted, but that these inputs also determine the state of the network. Thus, inputs from both the cerebellum and the cerebrum determine the probability of complex spike responses on a cycle-by-cycle basis providing a quasiperiodic framework to align synchronous groups. This sharply contrasts with a view in which the olive is a clock with regular periodicity. A circuit-wide understanding of cerebellar resonances on the basis of such a mechanism could open a novel pathway to explore the cerebellar gating by other brain regions.

The combination of delayed gap junctions and delayed inhibition, as found in the olivo-cerebellar loop (102, 110), can affect oscillatory behavior (111). The interplay between STOs and delayed inhibition is therefore also relevant for other neural circuitries, for instance for creating filter settings for the perception of sounds with specific oscillatory properties (112–114) or orchestrating rhythmic movements as shown in the present study (see also (115)).

### A ratchet rather than a metronome

Well-coordinated movement sequences are not timed rigidly; they must be enacted flexibly and contextually. In order to catch a ball, or a prey, or to perform any other appropriately timed movement, it is essential to fine-tune the duration and onsets of multiple coordinated output systems. An inferior olive that responds contextually to time varying input by advancing and delaying cycles does not act as a rigid clock or metronome, but more contextually, as a ratchet-pole system, with the frequency of ‘clicks’ of the ratchet reflecting the recent history of applied torque. The properties we have encountered in this study are consistent with a ‘ratchet-like’ dynamics for the inferior olive, which integrates time-varying stimulus in a phase-dependent manner. According to this view, the inferior olive responds to all inputs (sensory and otherwise), by producing phase changes that are informative about the recent history of input, and dictate the appearance of complex spike waves arriving at the cerebellar cortex.

## METHODS

### Ethics Statement

All experimental procedures were approved *a priori* by an independent animal ethical committee (DEC-Consult, Soest, The Netherlands) as required under Dutch law.

### Animals

Experiments were performed on 16 adult (9 males and 7 females of 25 ± 14 weeks old) homozygous Gjd2^tm1Kwi^ (Gjd2 KO) mice which were compared to 15 wild-type littermates (8 males and 7 females of 26 ± 13 weeks old; means ± sd). The generation of these mice has been described previously (116). The data described in S2 Fig. originated from previously published recordings in 35 wild-type mice (61). All mice had a C57BL6/J background. The mice received a magnetic pedestal that was attached to the skull above bregma using Optibond adhesive (Kerr Corporation, Orange, CA) and a craniotomy of the occipital bone above lobules crus 1 and crus 2. The surgery was performed under isoflurane anesthesia (2-4% V/V in O_2_). Post-surgical pain was treated with 5 mg/kg carprofen (“Rimadyl”, Pfizer, New York, NY) and 1 μg lidocaine (Braun, Meisingen, Germany). Mice were habituated during 2 daily sessions of 30-60 min.

### Electrophysiology

Extracellular recordings of Purkinje cells were made in the cerebellar lobules crus 1 and 2 of awake mice as described previously (28). Briefly, an 8 × 4 matrix of quartz-platinum electrodes (2-4 MΩ; Thomas Recording, Giessen, Germany) was used to make recordings that were amplified and digitized at 24 kHz using an RZ2 BioAmp processor (Tucker-Davis Technologies, Alachua, FL). The signals were analyzed offline with SpikeTrain (Neurasmus, Rotterdam, The Netherlands) using a digital band-pass filter (30-6,000 Hz). Complex spikes were recognized based on their waveform consisting of an initial spike followed by one or more spikelets. A recording was accepted as that of a single Purkinje cell when a discernible pause of at least 8 ms in simple spike firing followed the complex spikes and when the complex spikes were of similar shape and amplitude throughout the recording.

Sensory stimulation was applied as air puffs of 20 psi and 25 ms duration directed at the whisker pad ipsilateral to the side of recording. The stimuli were given in trains of 100 or 360 pulses either at regular or alternating intervals. During a recording, trains with different stimulus intervals were played in a random sequence.

### Whisker movement tracking

Whisker videos were made from above using a bright LED panel as backlight (λ = 640 nm) at a frame rate of 1,000 Hz (480 × 500 pixels using an A504k camera from Basler Vision Technologies, Ahrensburg, Germany). The whiskers were not trimmed or cut. Whisker movements were tracked offline as described previously (117) using a method based on the BIOTACT Whisker Tracking Tool (118). We used the average angle of all trackable large facial whiskers for further quantification of whisker behavior.

### Complex spike pattern analysis

Of each Purkinje cell we computed the probability density function (PDF) of both its complex spike autocorrelogram and its distribution of intervals between consecutive complex spikes (inter-complex spike intervals (ICSIs)). PDFs were calculated with an Epanechnikov kernel with a width of 10 ms. In order to exclude stimulus-induced alterations in complex spike firing, complex spikes detected between 20 and 200 ms after a stimulus were omitted from this analysis. PDFs were calculated from 0 up till 500 ms. The peak in the ICSI PDF was considered as the “preferred ICSI interval” and its strength was expressed as the Z-score by dividing the peak value by the standard deviation of the PDF. To avoid having Purkinje cell recordings with relatively low preference for specific ICSI intervals to distort our analysis we wanted to specifically also look at the Purkinje cells with a high Z-scores. Therefore, we grouped Purkinje cells into high and low level Z-scores, using a threshold of 3.

Air-puff stimulations frequently triggered double complex spike response peaks, suggestive of an underlying inferior olivary oscillation. For further analysis of the conditional responses, an estimate of the putative inferior olivary frequency was derived from the interval between these two response peaks. First, it was established for each Purkinje cell whether two peaks were present in the PSTH. To this end, we set a threshold for each of these two peaks. For the first peak, this was calculated by reshuffling the ICSIs over the recording followed by calculating a stimulus-triggered pseudo-PSTH, repeating this procedure 10,000 times and selecting the 99% upper-bound. We considered the first response peak to be significant if it crossed the upper-bound uninterruptedly for at least 10 ms. Since the second response peak typically is much smaller than the first one, we calculated a new threshold for the second peak by excluding the time-window for the first response peak. This window was set from the time of the stimulus until where the response probability drops to the average response frequency, the response frequency as expected if stimuli do not trigger complex spikes, following the significant ‘first’ responsive peak. In 5 out of 98 Purkinje cells, the PDF of the response rate between clear peaks remained above the average response rate, in which case we used the time point where the amplitude drop in the PDF was more than twice the difference between upper bound and average response probability. The rest of the bootstrap method was identical to that for the first response peak. Only peaks up to 0.5 s after the stimulus were included in the population analysis.

### Prediction of response probability based on inferred olivary oscillation frequency

In order to test whether the phase of the inferior olivary oscillations affected the complex spike response probability, we compared the complex spike intervals over an air-puff for each stimulus that triggered a complex spike. To this end, we analyzed the recordings of 25 Purkinje cells (10 WT and 15 Gjd2 KO) that were measured previously in crus 1 and crus 2 of awake, adult mice. We included only Purkinje cells that displayed clear oscillatory complex spike firing indicated by the display of a secondary complex spike response peak, as evaluated according to the bootstrap method described above, and/or significant peaks in the ICSI histogram. Only stable recordings covering at least 500 stimuli at frequencies below 1 Hz were considered for this analysis.

For each recording we compared two idealized statistical models of the observed ICSI distributions: an oscillatory model showing phase-dependent spiking and stable olivary oscillation frequencies and a uniform model lacking phase-dependencies. For the oscillatory model, we created complex spike probability functions for the pre-stimulus interval (−300 to 0 ms) based on the oscillatory period established either for the ICSI distribution or from the interval between the two complex spike response peaks. We fitted a sine wave with the observed frequency, having its peak at the moment of the first complex spike in the stimulus response window (20-200 ms after the air puff) and derived spike probability levels during the pre-stimulus interval from these fits, with the trough representing zero probability. Frequency and amplitude of every cycle were kept constant for the whole recording. In the uniform model, we calculated the pre-stimulus spiking probability with a uniform distribution based on the complex spike frequency of each Purkinje cell. We did include a refractory period, being the shortest ICSI observed for each recording, to reflect the inability of consecutive complex spikes to occur with a very short time interval. Refractory periods were comparable between mutants and wild types; 49 ± 15 ms for Gjd2 KO cells and 50 ± 20 ms for WT cells. Subsequently, we constructed compound fits consisting of linear summations of the two models. One extreme was the oscillatory model and the other the uniform model and we considered nine intermediate combinations (e.g., 0.3 × the oscillatory model + 0.7 × the uniform model). Every compound fit was run for 10,000 times. The goodness of fit was computed as the absolute differences of every single run of the model with the actual ICSI distribution.

### Data Availability

Summarized data and analyses scripts are available online at https://osf.io/6x5Uy/. README files are provided at the same repository.

### Biophysically plausible 10 model

The model networks used here are comprised of a topographical grid of 200 coupled cells, in a 10×10×2 lattice arrangement, which may resemble an area of about 400 μm × 400 μm of the inferior olive, for instance, the rostral portion of the dorsal lamella of the principle olive of the mouse. It is available online at https://github.com/MRIO/OliveTree. branch ‘Warnaar’. For instructions on how to run the model and reproduce analysis, check README_Warnaar.txt’.

### Single cell model

Each cell within these networks was modelled according to the single cell model described in (46), which is an elaboration of a previous model (62) with an added axon and modified fast sodium channel. Equations are provided in the appendix of that publication at (https://doi.org/10.1371/journal.pcbi.1002814.s002), and can be checked in the MATLAB functions *IOcell* and *createDefaultNeurons* in the codebase. The model includes three compartments (soma, dendrite, axon hillock) with 12 conductances. In addition to the ionic mechanisms, the dendrite of the model cell has a Ca^2+^ concentration state variable, which is related to the intrusion through the Ca_v_2.1 channels. The main ionic conductances responsible for the oscillation are the somatic T-type Ca^2+^ and the Ca^2+^-activated K^+^ (SK) channels present in the dendritic compartment. The crucial parameters governing the emergence of subthreshold oscillations are randomized, reflecting the experimental facts that about one third of the cells oscillate endemically (*in vivo)* with intrinsic variations in oscillatory frequencies (43). The behavior of the STO of the model cells in our network as a function of their parameters, for models with and without gap junctions, are included in S5 and S6 Fig. Cell parameters are found in supplementary table 1.

### Gap junction connectivity

Connectivity is created with the function ‘createW.m’ in the MATLAB codebase. Briefly, cells within a specified radius of each other were connected according to a probability function such as to ensure the specified mean degree in the network (n = 8), chosen to resemble the observed connectivity distributions reported in the literature. The connectivity parameters (distance and average connection probability) were chosen to match experimental values (radius ≤ 120 μm) and average connection probability (~8 neighbors). The procedure to obtain connectivity is as follows. First, pairwise distances between all cells are calculated. Then, a binary adjacency matrix is created by thresholding those distances within a specified radius. Thereafter, we assign a random number between 0 and 1 to each link from a uniform distribution. Finally, this matrix is made binary by comparing each entry with a probability so that the average number of connections approximates a given mean connectivity. This binary adjacency matrix is then multiplied by the mean gap junction conductance parameter. Finally, gap junction conductance values are then randomized by a uniform jittering of the conductance by 10% of their original value.

### Gap Junctions

The conductance of gap junctions was normalized with a saturating factor by difference of potential between the neighboring cells, according to (62) based on findings from (119) with the following function:

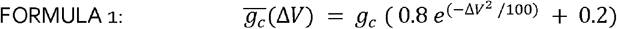

Where Δ*V* is the voltage difference between the connected cells, *g_c_* is the nominal coupling and *g_c_* is the effective coupling.

### Network inputs

Two inputs are given to the model, one emulating the sensory input from whisker pad stimulation and the other representing a stimulus-independent background reflecting diverse excitatory and inhibitory inputs to the inferior olive. The latter consisted of a continuous stochastic process with known mean and standard deviation with a relaxation parameter following the Ornstein-Uhlenbeck process (93), succinctly described underneath.

Only one subset of cells in the center of the network (40% the cells in a mask spanning a radius of 3 cells from the center of the network) representing efferent arborization, receives the “sensory input”, with “sensory” currents being delivered to the soma of modelled cells (g_AMPA_ = 0.15 mS/cm^2^). “Sensory input” was modeled according to O’Donnel etal. (120). The mask is represented in Fig. 5A.

The cells of the inferior olivary network most likely share input sources due to overlapping arborizations of efferent projections (121). To represent both shared and independent input, we have modeled the current source in each cell as having an independent process and a shared process, with a mix parameter (alpha) of 10% input correlation shared by all the cells in the network. This level of correlation leads to a coherent background oscillation in the cells of the network, which is exacerbated in the presence of gap junctions (S5 Fig.).

### Ornstein-Uhlenbeck noise process

Ornstein-Uhlenbeck (OU) is a noise process that ensures that the mean current delivered is well behaved and that the integral of delivered current over time converges to a constant value (93). The OU current is a good approximation for synaptic inputs originating in a large number of uncorrelated sources, where synaptic events are generated randomly and each event decays with a given rate (τ). We use a recursive implementation according to the following recursive formula:

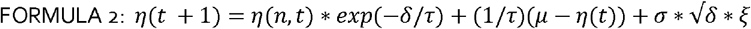

Where ⍰(*t*+1) is the noise value at *t*+1. The noise process is parameterized by *τ,σ,μ* where *τ* represents the synaptic decay time constant, *t* is the integration step time for our forward Euler integrator, *σ* is the standard deviation of the noise process and *μ* is its mean. The random draw from a Gaussian distribution at every time step is represented by r_i_.

Neurons in the inferior olive receive broad arborizations, signifying correlations in input sources across neurons. In our model we represent that via a mixture of an independent process for each neuron *n*_*independent*_ and a shared process, *n_all_*, common to all the neurons in the network, parameterized by a mixing parameter:

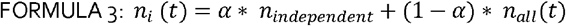

Simulation results throughout the article come from simulations with noise use an *α* where neurons share 10% of their noise input, though the main results are robust to changes in this value.

### Network Parameter Spaces

To examine the dependence of network dynamics on the characteristics of the incoming input, we computed the 200 neuron network sweeping a grid of the main input parameters (*τ,σ,μ,α*) of the Ornstein-Uhlenbeck noise process. The network response in terms of STO frequency, population firing rates, proportion of firing neurons was analyzed with respect to a grid of input parameters. For comparability of statistics and reproducibility of results, all results displayed in this article were obtained from a single random seed. We have tested the network with multiple seeds and the results are qualitatively indistinguishable.

The parameters of the Ornstein-Uhlenbeck process were tuned such that the network emulating the wildtype network (with gap junctions, WT) produced an average frequency of 1 Hz and more than 95% of the model cells fire at least once every 5 seconds (the parameter space for the network responses including STO, population firing rate and proportion of cells that fire within 3s is found in S4B Fig.). The parameters to achieve these criteria are dependent on the total leak through the gap junctions. There are multiple methods to compensate the absent leak in the gapless network. In the present case, the network without gap junctions has been tuned to produce the same firing frequency as the network with gaps by increasing the membrane leak currents from 0.010 to 0.013 mS/cm^2^. This results in a similar excitability but slightly lower STO frequency in the “mutant”. The average firing rate behavior of the network shows a linear relationship with the standard deviation of the OU process (S4 Fig.). For the present network with balanced connectivity and a single gap conductance of 0.04 mS/cm^2^, the Ornstein-Uhlenbeck parameters are *(τ,σ,μ,α),μ* = −0.6*pA*/*cm*^2^,*σ* = 0.6 *pA*/*cm*^2^ and *τ* = 20 *ms. τ* is a decay parameter that represents the synaptic decay times expected for olivary inputs, in this case chosen to emulate dendritic GABA according to Devor and Yarom (122).

### Synchrony and Frequency Estimation

Both synchrony and instantaneous frequency were estimated on the basis of a novel phase transformation of the membrane potential, which is more robust than the standard Hilbert transform, and can produce a linear phase response to the non-linear shape of the subthreshold oscillations (123). This transformation improves the estimation of the momentary phase and compensates for the fact that ionic mechanisms have different rates. To that end we have used the DAMOCO toolbox (124). From the instantaneous phase, the instantaneous frequency is simply the inverse of the first order finite difference of phases. Synchrony across cells is estimated with the Kuramoto order parameter:

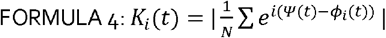

Where □i is the phase of each neuron, N is the number of neurons and ⍰ is the phase average of all oscillators.

### Post-Spike Phase Responses

To estimate the phase response curve of the stimulated neurons, first a “sensory stimulus” is delivered at a phase known to produce an action potential (and resetting). The location of the first peak after stimulation is recorded. Subsequently, eight more simulations receive another stimulus, with same parameters as the resetting stimulus, but at different phases (with an interval of 2⍰/8). The effect of that stimulation (delay or advance) on the next peak is recorded as a phase delta. Results are plotted in Fig. 7.

## Acknowledgements

The authors wish to thank llja IJpelaar and Mandy Rutteman for technical support, Sungho Hong, Marcel De Jeu, Jornt De Gruijl, Tycho Hoogland, Marcel van der Heide, Michiel Ten Brinke, Marylka Uusisaari, Yosef Yarom, Ben Torben-Nielsen and Rodolfo Llinás for memorable discussions, and nVidia for providing Tesla GPU cards on which the simulations were run.

## Author contributions

Conceptualization: M.N., P.W., L.W.J.B.; Data acquisition: M.N., P.W., V.R., C.B.O., S.L., E.I., L.W.J.B.; Network Modeling: M.N.; Formal analysis: M.N., P.W., V.R., C.B.O., S.L., E.I., L.W.J.B., C.I.D.Z.; Funding acquisition: C.I.D.Z.; Investigation M.N., P.W., V.R., C.B.O., L.W.J.B.; Methodology: M.N., P.W., L.W.J.B.; Project administration: M.N., C.I.D.Z.; Software: P.W., M.N.; Supervision: M.N., L.W.J.B., C.I.D.Z.; Visualization: P.W., M.N., L.W.J.B., C.I.D.Z.; Writing – Original draft preparation: M.N., P.W,.; Writing – Review & Editing: M.N., L.W.J.B., C.I.D.Z..

## SUPPLEMENTARY FIGURE CAPTIONS

**Fig. S1.**
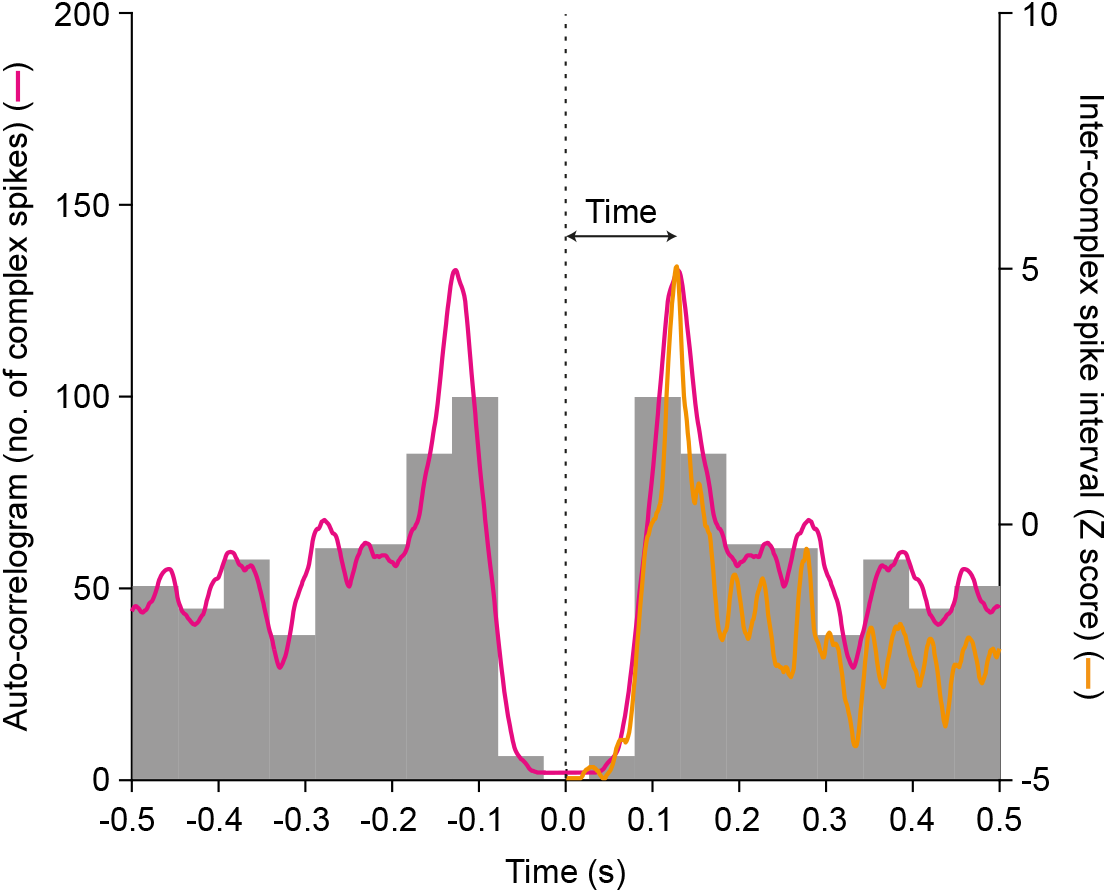
Comparison ofautocorrelogram with inter-complex spike interval histogram. The autocorrelogram of the same Purkinje cell as shown in Fig. 2A (10 ms bins) convolved with a 5 ms kernel (red curve) compared to the convolved inter-complex spike interval (ICSI) histogram. The auto-correlogram includes also intervals between non-consecutive complex spikes. Consequently, on a short time scale, both auto-correlogram and ICSI histogram are identical, but they diverge at longer time scales.

**Fig. S2.**
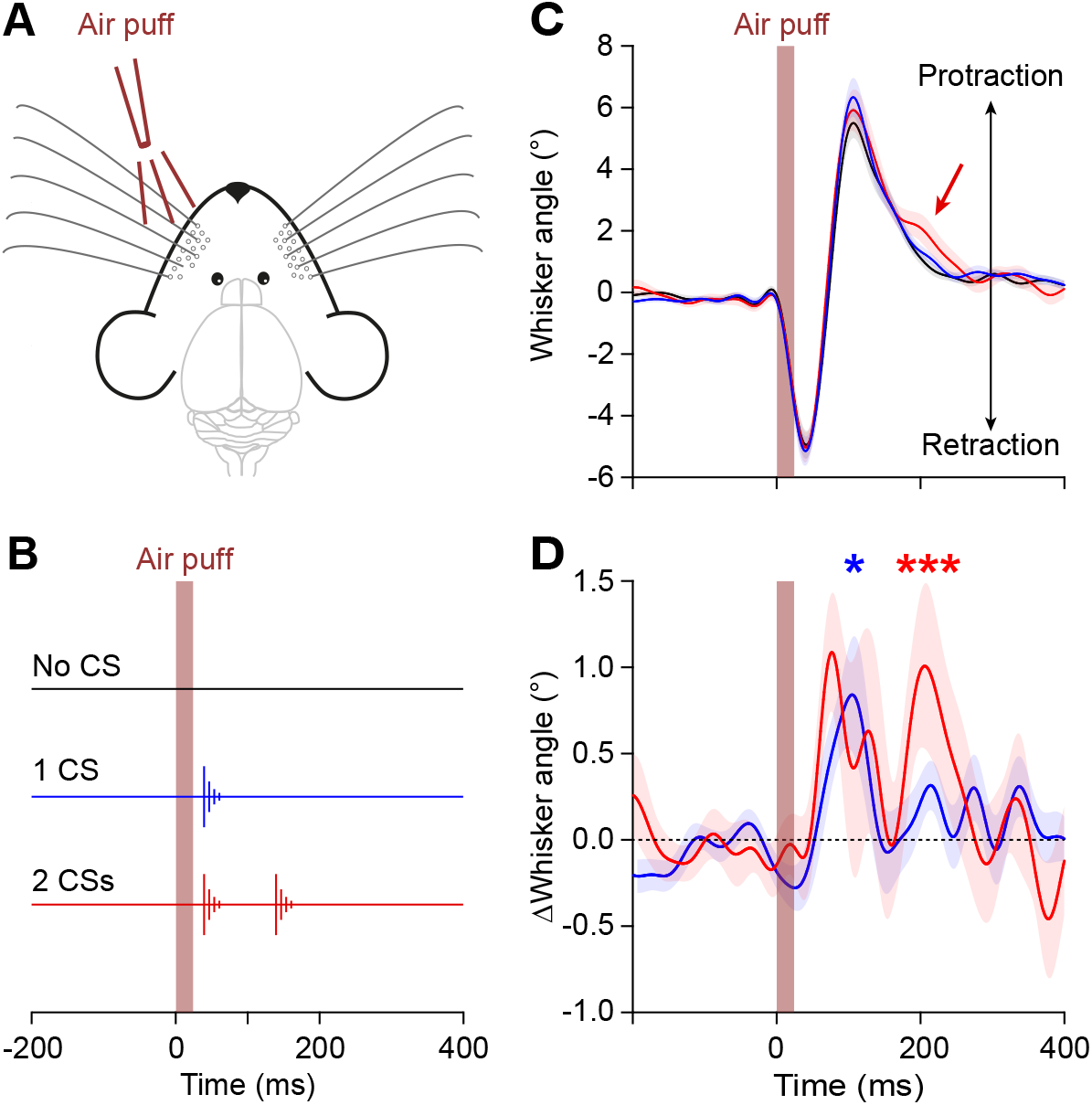
Complex spikes correlate with whisker protraction. (**A**) Mice received air puff stimulation of their whisker pad. (B) Purkinje cell recordings during sensory stimulation revealed that most trials had either no, one or tho complex spikes during the 200 ms after stimulus onset. We separated the trials according to this classification. Note that for the trials with a single complex spike, we considered the trials with a complex spike during the first 100 ms, but not during between 100 and 200 ms. (**C**) Averaged whisker traces (based on n = 35 Purkinje cells) show a reflexive whisker movement triggered by the air puff, consisting of an initial backward movement (largely caused directly by the air flow) followed by an active protraction. Trials in which a complex spike was detected only during the first 100 ms after the stimulus (blue line) had on average a slightly larger protraction than the trials without a complex spike (black line). The trials with two complex spikes also had a stronger protraction than the trials without a complex spike, but showed in addition a more protracted position later on during the trial (red arrow and red line). (**D**) Averaged subtracted traces showing the differences between trials with, respectively, a single complex spike (blue line) and two complex spikes (red line) and the trials without a complex spike. The occurrence of recurrent complex spike firing was thus reflected in the behavior of the mice. Shaded areas indicate the SEM. * *p* < 0.05; *** *p* < 0.001; Wilcoxon match-pairs test after Bonferroni correction for multiple comparisons.

**Fig. S3.**
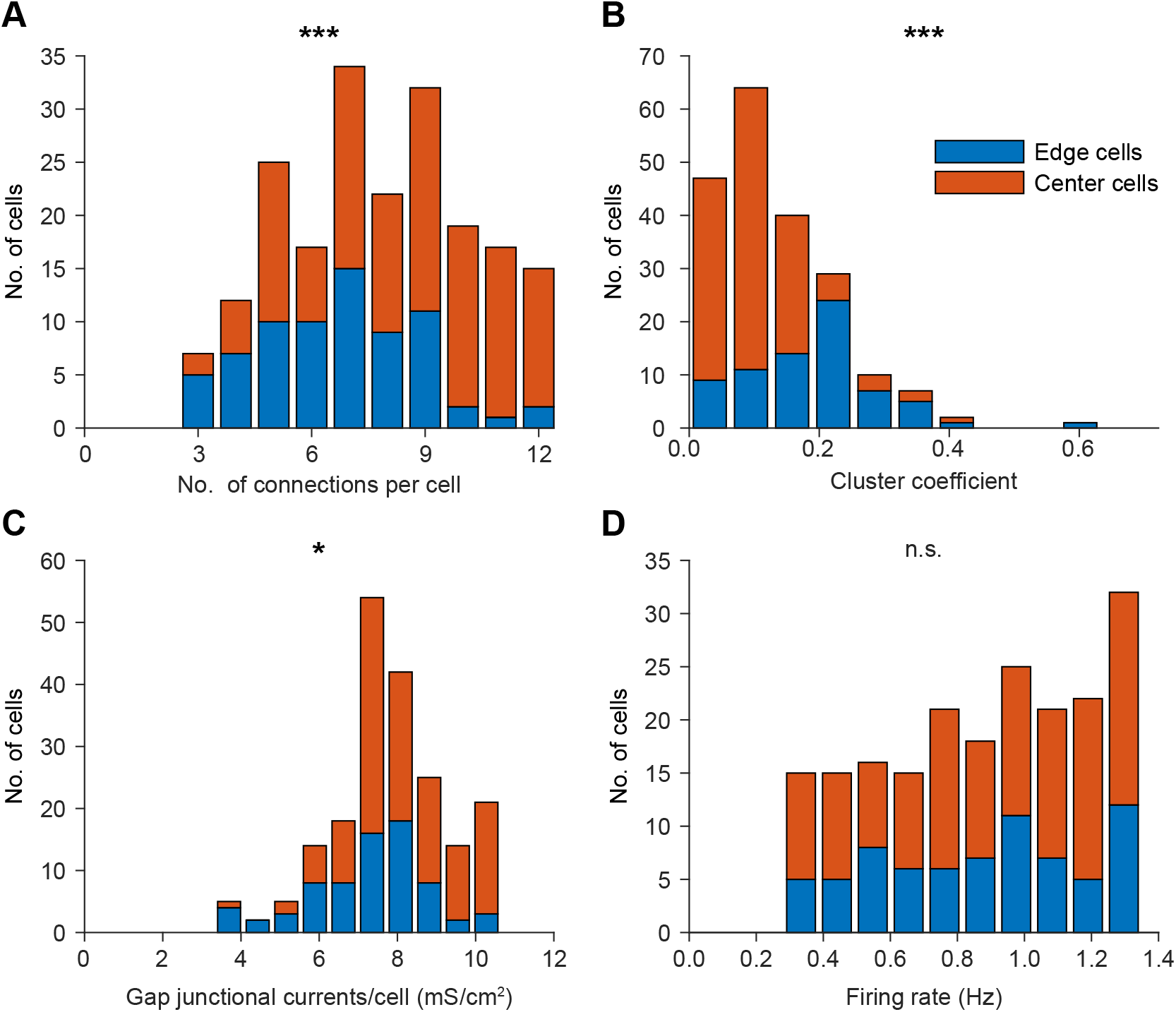
Behavior of cell on the edges of the model. The inferior olivary model has, as the inferior olive itself, boundaries. The impact of boundaries in connectivity of cells in the in vivo data (and by extension on the current leak through gap junctions) is, however, not known. The algorithm that generates connectivity enforces mean connectivity across cells, which increases the degree of clustering along the edges, but has at most a mild impact on the current leak through gap junctions. It is likely that the extra clustering degree along the edges may lead to a mild increase of coherence in STOs between neighbors, though this should not affect the overall conclusion - that the phase dependency of the STO under the presence of noise is at most short-lived. The data are represented as stacked bar plots. All data were tested using Kolmogorov-Smirnov tests.

**Fig. S4.**
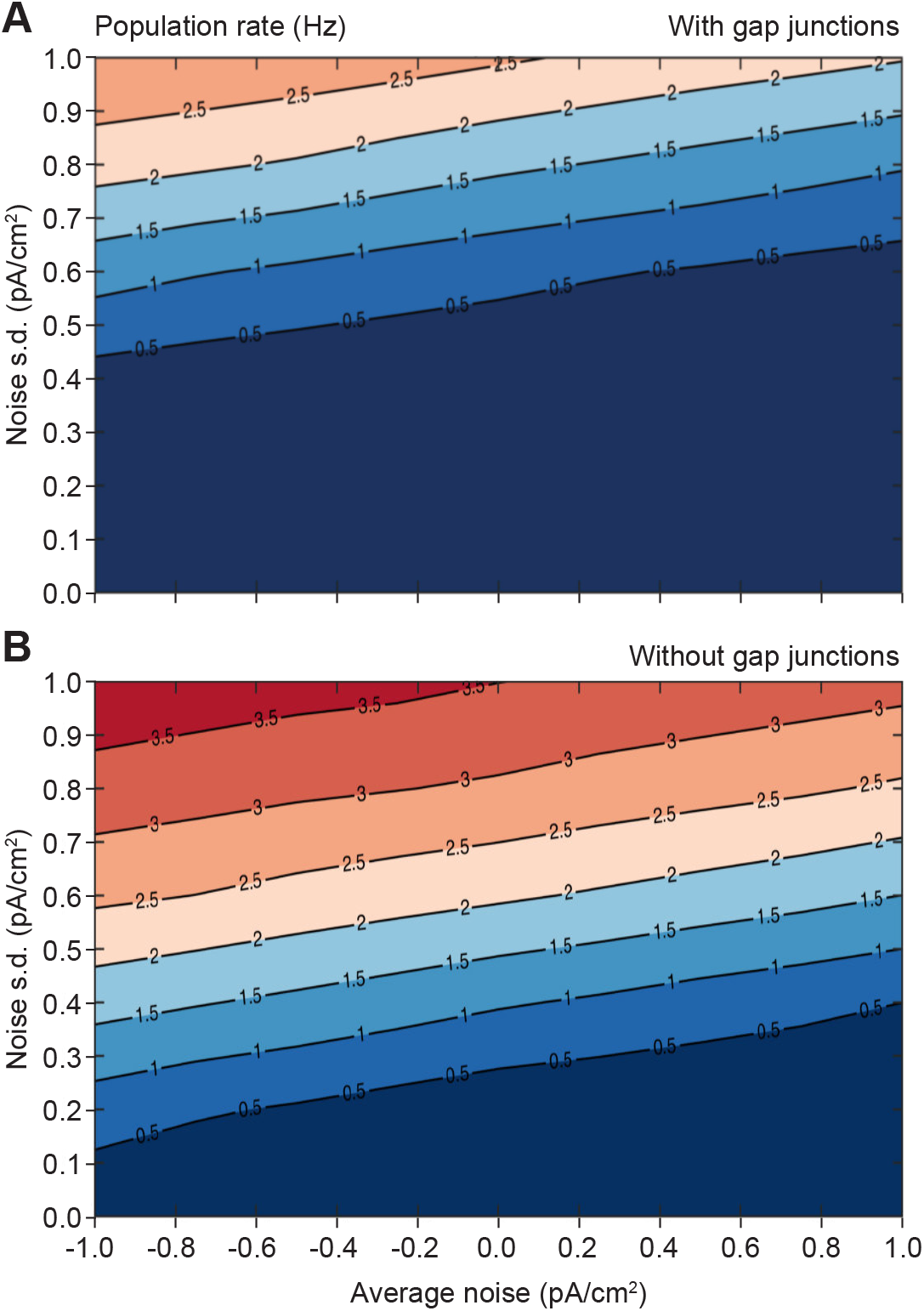
Contour plot indicating the average firing rate of networks as a function of the mean and standard deviation of the OU input. (**A**) Baseline input was chosen such that the cells of the model networks would produce approximately 1 Hz of spontaneous firing rate. The absence of gap junctional coupling in “mutant” model networks (**B**) leads to increased firing rate, which was compensated for by increasing the leak current of the membrane by 0.003 mS/cm^2^ (data not shown).

**Fig. S5.**
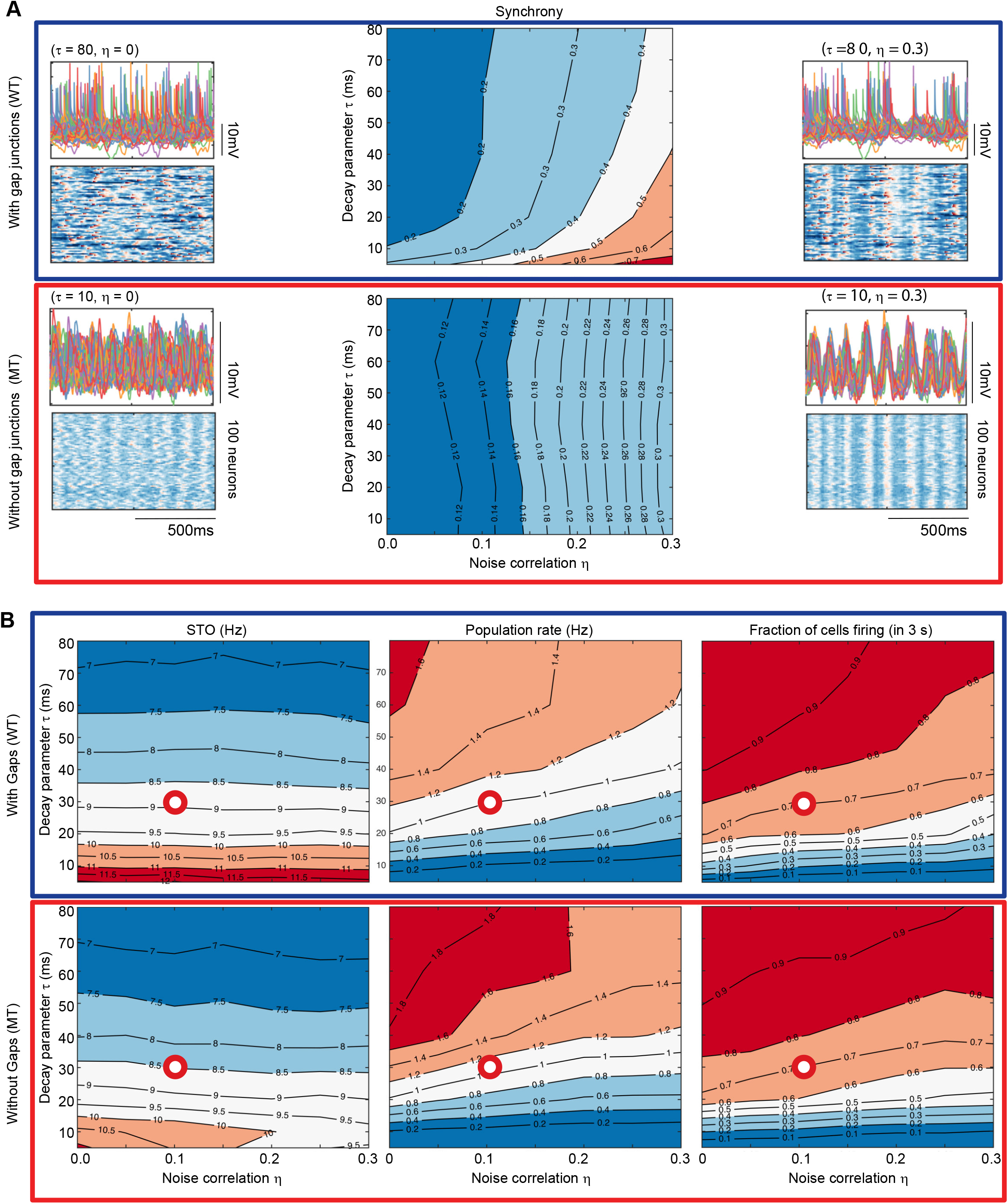
Behavior of model networks of identical composition as a function of the decay parameter of the Ornstein-Uhlenbeck process (τ) and the noise correlation (η). Model networks have been computed using a systematic exploration of the parameter space using 56 instances of the network model. The contour plots indicate isolines for synchrony (**A**), frequency of subthreshold oscillations (STOs), population firing rate and proportion of active cells (**B**). The results presented in the main text come from a model network with parameters chosen such as to display STOs with mean of 9 Hz, a population rate of about 1 Hz and with more than 70% of cells firing in three seconds (95% of cells fire within 10 s of simulation time). The position of this network in the contour plots is indicated with a red circle. In **A**, the thumbnails exemplify behaviors of extreme instances of the model network both as membrane potentials traces (top) and as heatmaps of the membrane potential (bottom). Arrows indicate parameter space coordinates of these examples. The decay of the Ornstein-Uhlenbeck process (τ) mostly impacts the firing rate of the model networks, while noise correlation (η) has a direct effect on synchrony. (**B**) Gap junctions amplify the input correlation given to the neurons, while having a minor effect on other aspects of the network dynamics.

**Fig. S6.**
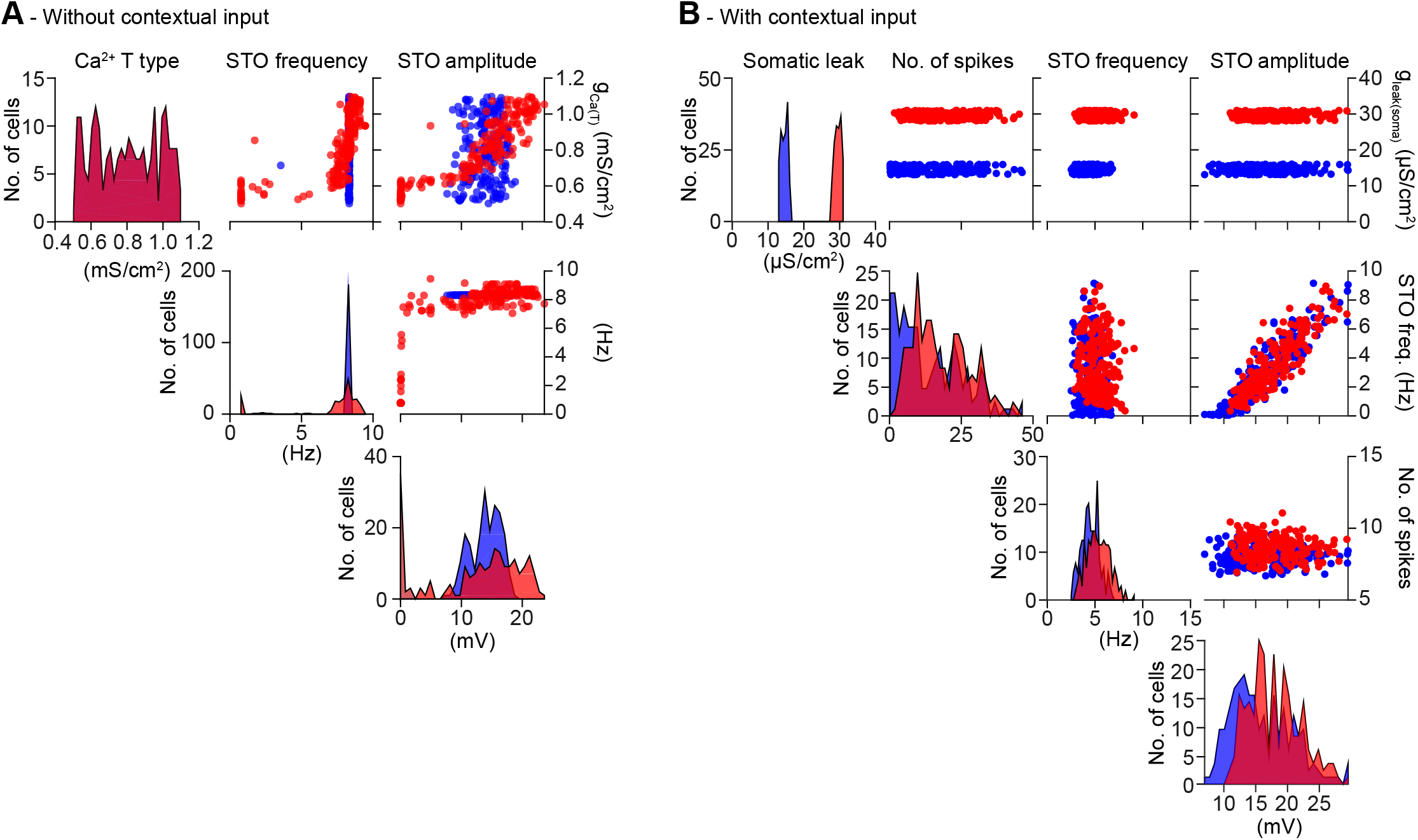
Physiological properties of the individual model cells as a function of Ca^2+^ T-type conductance in the absence of contextual input in the presence and absence of gap junctional coupling. (**A**) The Ca^2+^ T-type conductance is varied in the range of 0.5 to 1.1 mS/cm^2^, resulting in a range of oscillatory properties of the individual model cells. The left axis in the panels of the main diagonal display cell counts for the histograms. The right axis besides the rightmost panel displays the indicated continuous variable. The set of non-oscillating (zero amplitude, zero frequency) cells constituted about 25% of the model network in the absence of gap junctions (red), were engaged in the oscillation when gaps were added to these cells (blue). In the absence of contextual input the distribution of frequencies had peaks at zero and sharply synchronizes at approximately in the absence of Ornstein-Uhlenbeck input. (**B**) Comparison between the activity of networks with (WT)and without (MT) gap junctions under contextual input. MT cells have received compensation in the leak conductance of the cell. A slight increase of firing rate for the MT can be observed, and also a narrower distribution of frequencies, in comparison with the noiseless case.

**Fig. S7.**
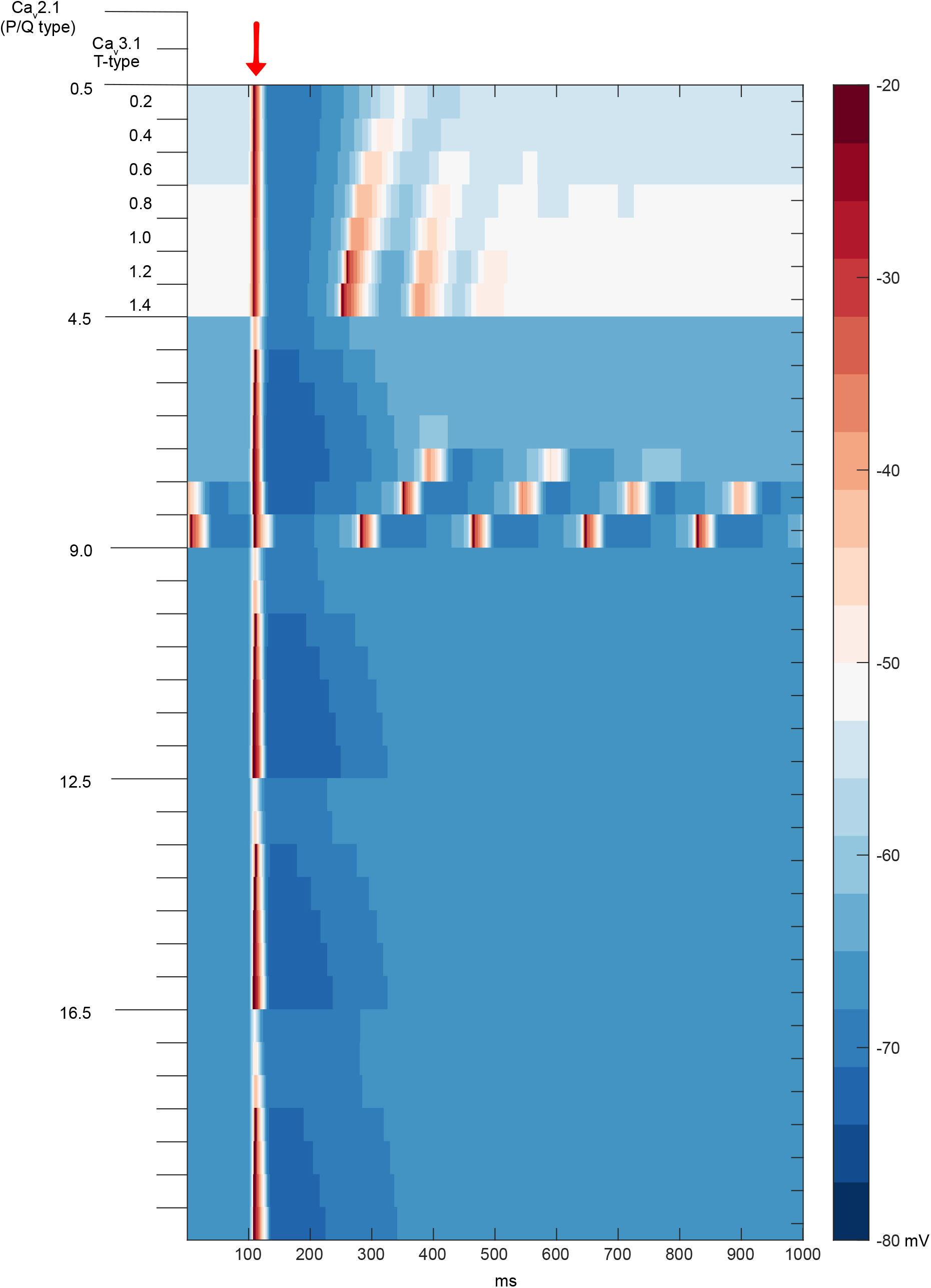
Rebound firing occurs only in a small region of parameter space. The post-spiking behavior of model neurons after a depolarization was examined for a grid of parameters, within known experimental values. A depolarizing current pulse triggers spikes at 100 ms. The rebound behavior is dependent on a complex interplay between a number of parameters, but most importantly, it exists in a rather narrow range of the parameter space between T- and P/Q-type Ca^2+^ channels. For very low levels of Ca_v_3.1 (T-type) expression, the model cell settles on an unphysiologically saturated depolarization after spiking. The cells in our model have Ca_v_2.1 (P/Q-type) expression around 4.5 and Ca_v_3.1 between 0.6 and 1.1.

**Fig. S8.**
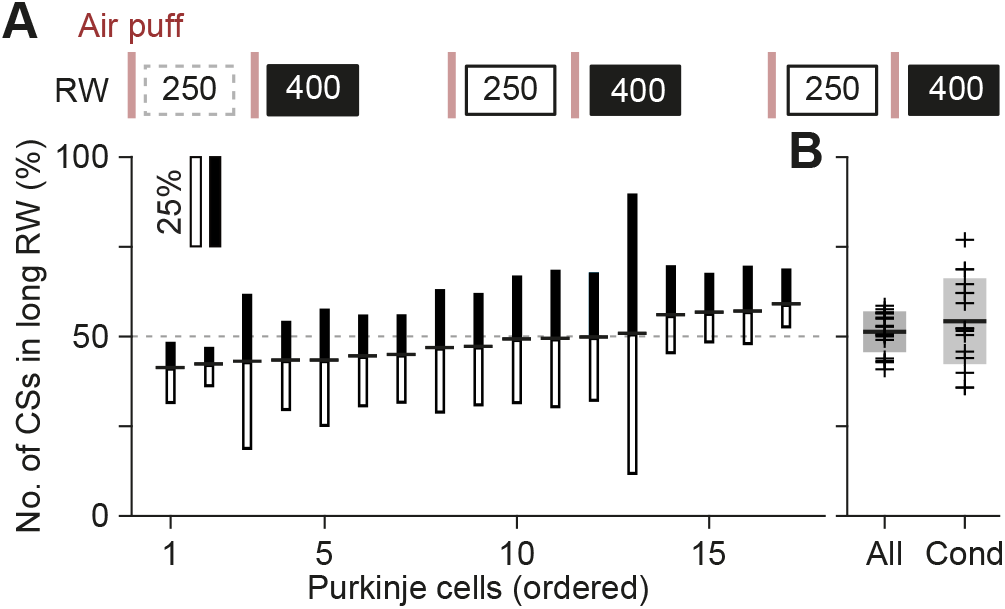
No phase-dependent spiking probabilities observed *in vivo*. (**A**) In order to test the impact of the presumed phase of the inferior olivary neurons in vivo, we applied a “gallop” stimulation pattern, alternating short (250 ms) and long (400 ms) intervals. Air puffs (vertical bars) were delivered to the whisker pad. Complex spikes were counted in response windows (RW) 20-200 ms post-stimulus and cells were sorted as a function of the ratio between the numbers of complex spikes in short and long intervals (indicated as horizontal dash between filled and empty bars). In this analysis, we included all RWs, irrespective of whether the preceding RW contained a complex spike or not (cf. Fig. 10). For each Purkinje cell the relative response probabilities for the long and short intervals are illustrated as the length of the filled and open bars, respectively. None of the Purkinje cells showed a significant difference in the response probability between the two intervals (all *p >* 0.05 on Fisher’s exact test). (B) Comparison of the response biases between all trials (“All”, **A**) and only those trials that followed an RW with at least one complex spike (“Cond”, Fig. 10A). Shown are the 25 and 75% quantiles and median per population.

