## supplemental table 1 for "Quasiperiodic rhythms of the inferior olive"

| **Parameter** | **fixed** | **range** | **unit** | **Notes** |
| --- | --- | --- | --- | --- |
| **Conductances** |  |  |  |  |
| **Soma** |  |  |  |  |
| Ca T-type (v3.1) |  | 0.6-1.1 | mS/cm^2^ | 1/3 oscillatory cells |
| Na | 150 |  | mS/cm^2^ |  |
| Kdr | 9 |  | mS/cm^2^ |  |
| K | 5 |  | mS/cm^2^ |  |
| leak | 0.016 |  | mS/cm^2^ | -0.003 for Cx36KO |
| coupling soma - dendrite | 0.13 |  | mS/cm^2^ |  |
| **Dendrite** |  |  |  |  |
| Ca act K |  | 35-45 | mS/cm^2^ | for ADHD variation |
| Ca H (P/Q) (v2.1) | 4.5 |  | mS/cm^2^ |  |
| leak | 0.016 |  | mS/cm^2^ |  |
| HCN |  | .12-1.12 | mS/cm^2^ | for ADHD variation |
| coupling dendrite - soma | .13 |  | mS/cm^2^ |  |
| **Axon** |  |  |  |  |
| Na | 240 |  | mS/cm^2^ |  |
| K | 240 |  | mS/cm^2^ |  |
| leak | 0.016 |  | mS/cm^2^ |  |
| coupling axon - soma | .13 |  | mS/cm^2^ |  |
| **Reversal Potentials (mV)** |  |  |  |  |
| Na | 55 |  | mS/cm^2^ |  |
| K | -75 |  | mS/cm^2^ |  |
| Ca | 120 |  | mS/cm^2^ |  |
| HCN | -43 |  | mS/cm^2^ |  |
| leak | 10 |  | mS/cm^2^ |  |
| **Capacitance** |  |  |  |  |
|  | 1 |  | pF/cm^2^ |  |
| **Synaptic Conductances** |  |  |  |  |
| GABA dendrite | 0.25 |  | mS/cm^2^ |  |
| GABA soma | 0.5 |  | mS/cm^2^ |  |
| AMPA dendrite | 1 |  | mS/cm^2^ |  |
| **Surface ratios** |  |  |  | all currents are normalized |
| soma/dendrite | 0.25 |  |  | by surface area |
| axon hillock / soma | 0.15 |  |  |  |
| **Reversal Potentials** |  |  |  |  |
| Na | 55 |  | mV |  |
| K | -75 |  | mV |  |
| Ca | 120 |  | mV |  |
| h | -43 |  | mV |  |
| leak | 10 |  | mV |  |
| GABA soma | -63 |  | mV | Devor and Yarom, 2002 |
| GABA dendrite | -70 |  | mV | Devor and Yarom, 2002 |
| AMPA | 0 |  | mV | O'Donnel et al., 2010 |
